# The Single-Stranded DNA-Binding Factor SUB1/PC4 Alleviates Replication Stress at Telomeres and is a Vulnerability of ALT Cancer Cells

**DOI:** 10.1101/2024.10.22.619621

**Authors:** Jean-Christophe Dubois, Erin Bonnell, Julie Frion, Samuel Zimmer, Muhammad Riaz Khan, Gabriela M. Teplitz, Lisa Casimir, Élie Méthot, Amélie Filion, Mouhamed Idrissou, Pierre-Étienne Jacques, Raymund J. Wellinger, Alexandre Maréchal

## Abstract

To achieve replicative immortality, cancer cells must activate telomere maintenance mechanisms. In 10-15% of cancers, this is enabled by recombination-based alternative lengthening of telomeres pathways (ALT). ALT cells display several hallmarks including heterogeneous telomere length, extrachromosomal telomeric repeats and ALT-associated PML bodies. ALT cells also have high telomeric replication stress (RS) enhanced by fork-stalling structures (R-loops, G4s) and altered chromatin states. In ALT cells, telomeric RS promotes telomere elongation but above a certain threshold becomes detrimental to cell survival. Manipulating RS at telomeres has thus been proposed as a therapeutic strategy against ALT cancers.

Through analysis of genome-wide CRISPR fitness screens, we identified ALT-specific vulnerabilities and describe here our characterization of the roles of SUB1, a ssDNA-binding protein, as a novel regulator of telomere stability. SUB1 depletion further increases RS at ALT telomeres, profoundly impairing ALT cell growth without impacting telomerase-positive cancer cells. During RS, SUB1 is recruited to stalled forks and ALT telomeres via its ssDNA-binding domain. This recruitment is potentiated by RPA depletion, suggesting that these factors may compete for ssDNA. The viability of ALT cells and their resilience towards RS also requires ssDNA-binding by SUB1. SUB1 depletion accelerates cell death induced by FANCM depletion, triggering unsustainable levels of telomeric damage specifically in ALT cells. Finally, combining SUB1 depletion with RS-inducing drugs rapidly induces replication catastrophe in ALT cells. Altogether, our work identifies SUB1 as a new ALT susceptibility with important roles in the mitigation of RS at ALT telomeres and suggests new therapeutic strategies for a host of still poorly managed cancers.

**Significance Statement:** Currently, there are few treatment options for ALT cancers with chemotherapy still occupying center stage despite often limited efficacy. ALT cancer cells experience high levels of replication stress at telomeres and its enhancement (e.g. via ATR inhibition) is a promising therapeutic strategy. Sensitivity to ATR inhibition varies amongst ALT cell lines/tumors warranting the development of additional ways to modulate telomeric replication stress. Here we identify SUB1, a single-stranded DNA-binding protein, as a vulnerability of ALT cells. SUB1 localizes to ALT telomeres and mitigates deleterious replication stress. SUB1 depletion synergizes with ATR inhibition and FANCM downregulation suggesting that co-targeting SUB1 with other regulators of replication stress at telomeres may kill ALT cancer cells more effectively.

## Introduction

Telomeres are transcriptionally active heterochromatic nucleoprotein structures that protect chromosome ends from nucleolytic degradation, illegitimate recombination and recognition by the DNA damage response (1). Human telomeres are composed of repetitive microsatellite DNA sequences (TTAGGG, ∼3-15 kb) with a 3’ G-rich overhang of 50-200 nts that folds back onto itself forming a structure termed t-loop. They are bound by shelterin proteins (POT1, TPP1, TRF1, TIN2, TRF2 and RAP1/TERF2IP) that are critical for telomere stability. Telomeres pose a particular challenge to the replication machinery due to their G-rich nature, which may support alternative DNA structures such as G-quadruplexes (G4s). Telomeres also express the TERRA long non-coding RNA which can form RNA:DNA hybrids at chromosome ends. Such structures slow down and stall ongoing replication forks, creating a state of DNA replication stress (RS) (2–4). Moreover, telomeres are origin-poor genomic regions, making fork stalling events more difficult to resolve. Consequently, RS frequently causes breaks in telomeric DNA which contribute to genome instability in cancer cells (5, 6).

Because of nucleolytic processing and the fact that DNA polymerases are unable to fully replicate chromosome ends, telomeres in somatic cells tend to lose 50-250 bp/cell division. Critically short telomeres induce an irreversible DNA damage signal that triggers senescence and cell death. This barrier to tumorigenesis is overcome by most cancers via the reactivation of telomerase (TEL+ cells). However, 10-15 % of cancers remain telomerase-negative and rely on alternative lengthening of telomeres pathways (ALT). ALT cells use homology-directed break-induced replication to elongate telomeres and to achieve replicative immortality (7, 8). ALT is particularly prevalent in cancers of mesenchymal origin (e.g. soft tissue sarcomas, osteosarcoma, glioma) and ALT tumours have poor prognoses due to complex karyotypes and a dearth of targeted therapies (4, 9). Cells maintaining telomeres with ALT have very long and heterogeneous telomeric repeat tracts that contain single-stranded regions, harbour extrachromosomal telomeric DNA called C-circles and their telomeres associate in subnuclear structures called ALT-associated promyelocytic leukaemia bodies (APBs). Mutation and/or protein loss of the ATRX/DAXX histone H3.3 chaperones is also highly prevalent in ALT cells (9, 10). Furthermore, telomeres in ALT cells experience higher levels of spontaneous RS and DNA damage than that observed in TEL+ cells (11).

The genotoxicity of RS both genome-wide and at telomeres is normally prevented by the DNA damage response (12). At stalled RFs with exposed single-stranded (ss)DNA, the DDR is initiated by the rapid binding of the RPA complex (RPA1,2,3). RPA quells secondary structure formation within exposed ssDNA and protects it from nucleolytic degradation. It also sequentially recruits and activates genome guardians including the checkpoint kinase ATR to signal RS, promote repair and restart stalled forks (13). RPA-ssDNA is particularly abundant at ALT telomeres which are under a quasi-permanent state of RS that needs to be appropriately managed to maintain chromosome ends (14–17). Importantly, RPA prevents the accumulation of unprotected ssDNA at telomeres and its depletion enhances telomere aggregation specifically in ALT cells (18). In addition to RPA, numerous fork remodeling enzymes and homologous recombination factors such as FANCM, SMARCAL1 and BLM were all shown to be important to achieve a proper level of DNA RS at telomeres compatible with ALT cell proliferation (19–28). Exacerbation of telomeric RS in ALT cells, for instance via ATR inhibition (ATRi), has indeed been gaining traction as a potential therapeutic strategy for these often hard-to-treat cancers (4, 14, 16, 17, 29–31).

In an effort to identify susceptibilities of ALT cancer cells, we analyzed large-scale CRISPR-based fitness screens (32, 33). Here, we characterize the evolutionarily conserved ssDNA-binding protein SUB1/PC4 as an ALT-specific vulnerability. SUB1 associates with ssDNA at stalled forks and damaged telomeres to resolve RS. In the absence of SUB1, telomeres become enlarged and accumulate damage, leading to decreased ALT cell viability. Co-depletion of FANCM and SUB1 further enhances telomeric RS to unsustainable levels specifically in ALT contexts resulting in a strong decrease in viability. Mechanistically, we show that SUB1 binds ssDNA to alleviate RS and protects against replication catastrophe induced by RPA depletion and/or ATRi/HU treatment. Accordingly, SUB1 depletion in ALT cells sensitizes to ATR or PARP inhibition, suggesting novel therapeutic strategies for ALT cancer cells.

## Results

### SUB1/PC4 is a Vulnerability of ALT Cancer Cells

To identify novel vulnerabilities of ALT cancer cells, we mined the Project Score and Achilles large scale cancer dependency datasets from the Wellcome Sanger and Broad Institutes (33, 34). Differential genetic dependencies were assessed across >18,000 genes for 7 experimentally validated ALT cell lines (U2OS, CAL72, CAL78, SKNFI, G292 clone A141B1, SAOS2 and HUO9) and compared with those of the remaining 801 cell lines (Depmap 21Q1, Table S1). Twelve statistically significant putative ALT-specific genetic dependencies were discovered using this strategy, and similar results were also obtained using raw CERES scores (Fig.1A,B, Fig. S1A,B). We note that a congruent set of ALT-specific vulnerabilities was recently found using a similar approach while our manuscript was under preparation (35). Interestingly, FANCM and its binding partners FAAP24 and MHF1/2, previously found as essential for sustained ALT cell proliferation, were all identified as top ALT-specific vulnerabilities, validating the robustness of this approach (24–26). Further in agreement with prior work, depleting Fanconi anemia core complex proteins FANCA, FANCF and FANCG appeared particularly impactful towards the ALT cell lines subset (26, 36). The SMARCAL1 DNA translocase, which mitigates DNA RS at telomeres also came up as a differential dependency of ALT cells (23, 28, 30).

**Figure 1.**
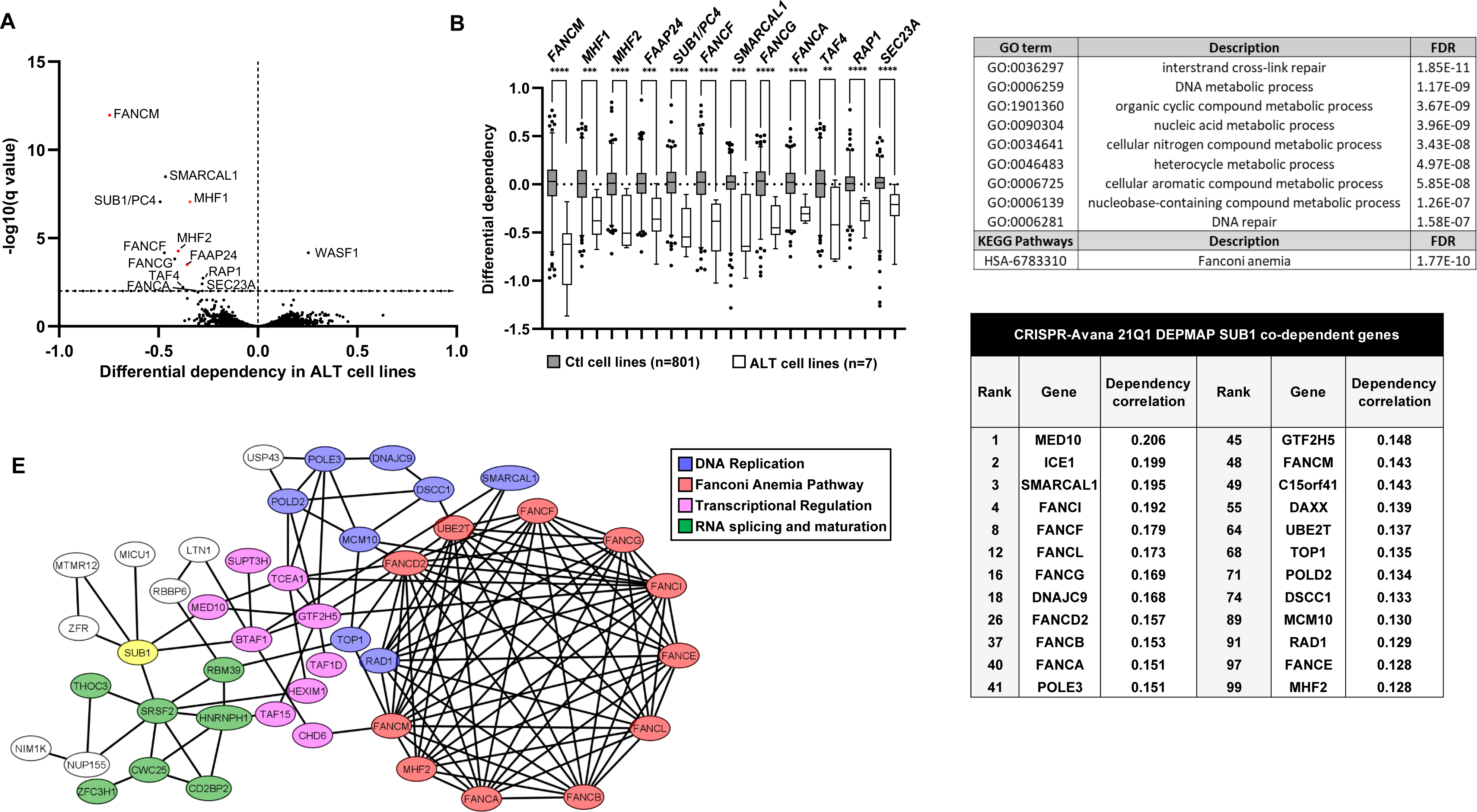
Identification of ALT-specific Vulnerabilities. **(A)** Volcano plot of the differential genetic dependencies in experimentally validated ALT cell lines. Mean subtracted CERES scores obtained from the Cancer Dependency Map (21Q1) for a panel of validated ALT cell lines (n=7) were compared with the rest of the available cell lines (n=801). Differential dependencies are plotted against the FDR (false discovery rate)-corrected Q value, calculated from multiple unpaired t tests. The dashed horizontal line represents an FDR of 1%. Red dots represent components of the FANCM complex. **(B)** Box and whiskers plots of preferentially essential genes of ALT cells. Horizontal lines represent the median and whiskers show 1-99 percentile cell lines. Significance was determined by Mann-Whitney tests. **** P<0.0001, *** P<0.001, ** P<0.01. **(C,D)** Co-dependency analysis of SUB1/PC4 (Depmap 21Q1) reveals association with the Fanconi Anemia DNA repair pathway and nucleic acid metabolism. **(E)** Interactome of top 100 SUB1/PC4 co-dependent genes created using STRINGdb (medium confidence setting (0.4)) and Cytoscape for network formatting.

Intriguingly, SUB1/PC4, a single-stranded nucleic acid-binding protein that functions as a transcriptional cofactor and a regulator of genome stability (37–41), displayed one of the strongest putative dependency of the ALT cells subset (Fig. 1A,B). Since many ALT cell lines in the Depmap are osteosarcomas, we further validated that SUB1 is particularly important for ALT compared to non-ALT osteosarcomas (Fig S1C, Table S2). Using the top 5% FANCM- or FANCM complex-(FANCM-FAAP24-MHF1/2) dependent cells lines in the Depmap also revealed SUB1 as a vulnerability, supporting the idea that SUB1 is important for FANCM-dependent cells (Fig. S1D,E, Tables S3,4). ALT cells were among the top SUB1-dependent cell lines with 5 out of 7 ALT lines used in the initial screening found within the top 20 SUB1-dependent lines (Table S5). As ATRX and DAXX mutations or deletions are frequently found in ALT cell lines, we also determined whether damaging point mutations in this histone chaperone complex were enriched in SUB1-dependent cells. Even though none of the validated ALT cell lines used for the initial differential dependency analysis harbored point mutations, at least 3 were previously shown to be completely devoid of ATRX expression and 12.5 % of the top SUB1-dependent cell lines (Top 2%, differential dependency <-0.4, 4/32) contained damaging ATRX or DAXX point mutations, a significant over-representation over the complete Depmap Avana CRISPR cell line compendium (4.5 %, 36/808 (21Q1)) (hypergeometric test P=0.038) suggesting that ALT characteristics correlate with vulnerability to loss of SUB1/PC4 (Table S6 and (10, 42)). Finally, a recent effort to analyze telomere maintenance using high-throughput DNA sequencing data from the Cancer Cell Line Encyclopedia and the Genomics of Drug Sensitivity in Cancer project identified 42 potentially ALT cancer cell lines and 24 of these had available CRISPR fitness screen data ((43) and Table S7). Analysis of differential dependencies across this larger panel confirmed the ALT-specific SUB1/PC4 differential dependency (Fig. S1F and Table S7). Co-dependency analyses showed that the impact of SUB1 depletion on cancer cell lines is most similar to that of transcriptional regulators ICE1 and MED10 but also to a large group of genome maintenance factors which prominently feature Fanconi anemia factors in agreement with its proposed roles at the interface between transcription and genome stability (Fig. 1C-E, Table S8 and (37)). Reanalysis of co-dependencies using the most recent Depmap version (23Q2) confirmed the close similarity of the impact of Fanconi factors and SUB1 depletion on an expanded set of ALT cell lines (Table S8).

### SUB1/PC4 Associates with Stressed Telomeres in ALT Cells

SUB1/PC4 is a small (15 kDa), evolutionarily conserved single-stranded nucleic acid binding protein that was shown to play important, but poorly understood roles in genome maintenance (40, 41, 44, 45). It has a bimodal domain architecture composed of unstructured N-terminal serine and lysine-rich domains and a C-terminal ssDNA-binding domain that forms dimers (Fig.2A and (46)). Amongst the ALT-specific dependencies that we identified on the Depmap, many were previously found to localize to ALT telomeres (SMARCAL1, FANCM, FAAP24 and RAP1 (23–27). To determine whether SUB1 also associates with telomeres in ALT cells, we performed immunofluorescence combined with telomere-specific *in situ* hybridization (FISH). Although RAD52, BLM or RPA spontaneously colocalized with telomeric foci in U2OS cells (Fig. S2A) as previously described (47, 48), we were unable to detect endogenous SUB1 foci at telomeres in unstressed ALT or non-ALT (HeLa LT) cells. Interestingly, SUB1 was recently found in the human replicating telomere proteome by qTIP-iPOND (49) suggesting that it may be present at telomeres in unstressed cells but at levels below the detection threshold of immunofluorescence. Prior work supports a clear contribution of ATR to telomere maintenance and stability in both ALT and telomerase-positive (TEL+) cells (5, 16, 50, 51) .To determine whether enhancing RS at telomeres would induce the recruitment of SUB1 to telomeres, we exposed cells to ATRi and performed FISH-IF. Indeed, ATRi treatment induced telomeric SUB1 foci formation in a substantial portion of U2OS cells (Fig. 2B-E). Enhancing the availability of ssDNA via RPA70 depletion further increased the association of SUB1 with telomeres (Fig. 2B-D) and also more generally to stalled replication forks (Fig. 5, see later part of the text). In stark contrast, combined depletion of RPA70 and ATR inhibition in TEL+ HeLa LT cells induced only marginal recruitment of SUB1 to telomeres Fig. 2E, S2B). Cell synchronization and quantitative image-based cytometry showed that most telomeric SUB1 foci formed during S-phase with a few foci being visible in G2 cells as well. Very little foci were detected in G1 cells, indicating that SUB1 associates with telomeres specifically when they encounter replication problems (Fig. 2F). The recruitment of SUB1 to telomeres depended on its ssDNA-binding capability as single (W89A) or triple (F77A/K78G/K80G) mutations (3XMut) within its C-terminus harboring the nucleic acid-binding domain (52) completely abrogated telomeric SUB1 foci formation (Fig. 2G, S2C,D).

**Figure 2.**
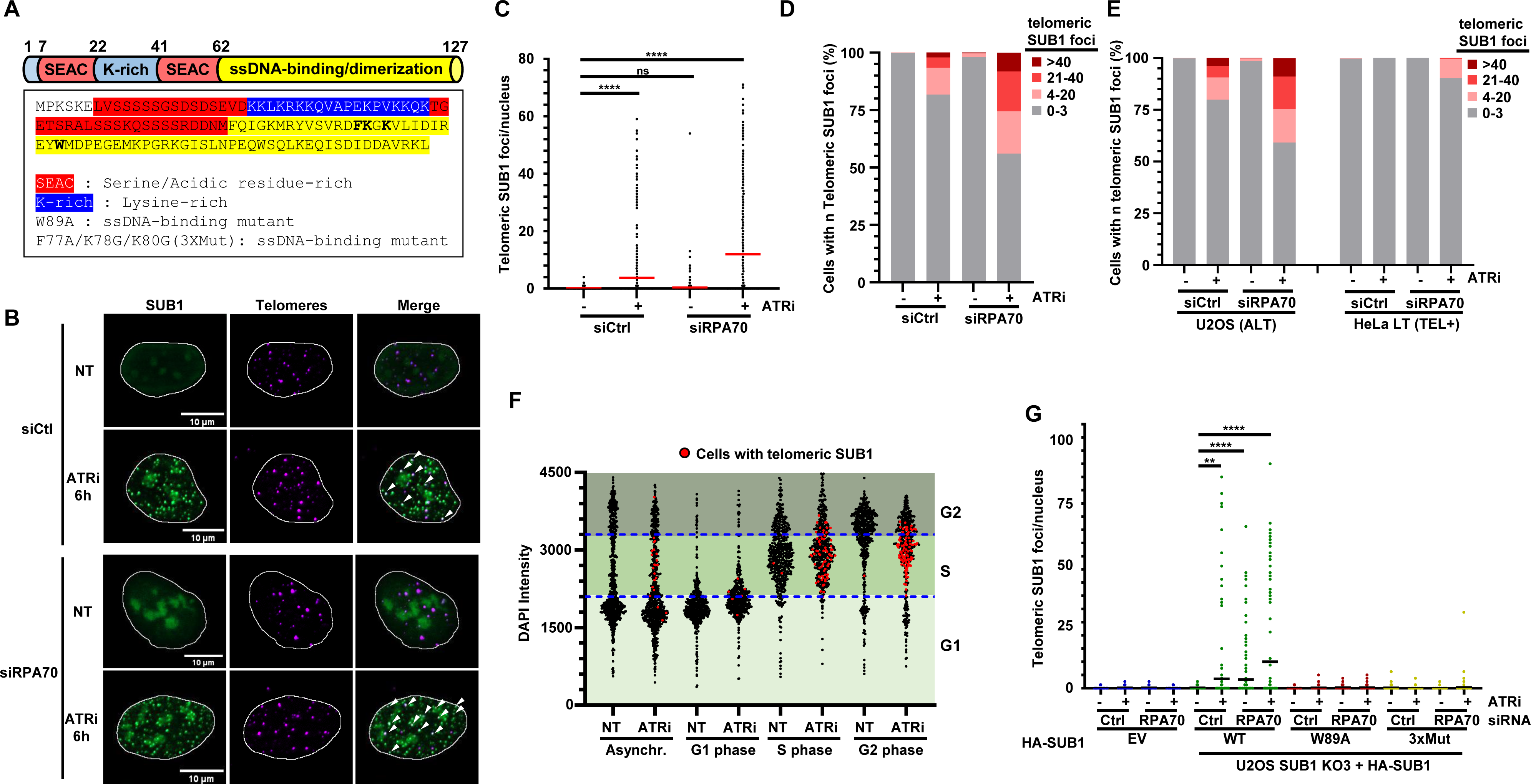
SUB1 localizes to damaged telomeres. **(A)** Rod representation and sequence of the different domains of SUB1 and position of ssDNA-binding domain mutations. **(B-E)** U2OS and HeLa LT cells transfected with indicated siRNAs were exposed to 10 μM ATRi (AZD6738) for 6h and processed for immunofluorescence. Telomeric SUB1 foci were automatically counted using CellProfiler. (**B)** Representative U2OS cells are shown. White arrows show colocalization events. **(C)** Graphical representation of the number of telomeric SUB1 foci per nuclei. Each dot represents one nucleus and the red line is the mean number of foci/nucleus for each condition. Graph represents the merged data from two biological replicates. Statistical significance was established by Ordinary one-way ANOVA and Dunnett’s multiple comparison tests. **(D-E)** Histogram representing the percentage of cells with 0-3, 4-20, 21-40 or >40 telomeric SUB1 foci in U2OS or HeLa cells. Graph represents the merged data from two biological replicates. **(F)** U2OS cells synchronized by double thymidine block were exposed to ATRi for 3 hrs prior to TEL-FISH and DAPI staining. Quantitative image-based cytometry was performed to assess cell cycle distribution of cells containing ≥ 4 telomeric SUB1 foci. **(G)** Scatterplot representing the number of SUB1 telomeric foci in KO SUB1 cells (KO3) complemented with WT or mutant HA-SUB1. The SUB1 KO3 cell line is described in more details in figures 3 and S3. The graph represents the merged data from two biological replicates. Statistical significance was established by Kruskal-Wallis and Dunn’s multiple comparison tests (P<0.05 *, P<0.01 **, P<0.001 *** and P<0.0001 ****).

#### SUB1 Alleviates RS at Telomeres in ALT Cells

As SUB1 associates with stressed telomeres, we examined whether it could be involved in telomeric RS. Transient depletion of SUB1 in a panel of ALT+ (3) and TEL+ (3) cancer cell lines induced an accumulation of RPA32 pS33 exclusively at ALT telomeres (Fig. 3A, B and S3A). The RS induced by SUB1 depletion also occurred outside of telomeres as about 50 % of RPA32 pS33 foci did not co-localize with Tel-FISH signal (Fig. 3A). These phenotypes were not observed in TEL+ cells (Fig.3A,B). The BLM helicase also formed bright punctate foci colocalizing with telomeres upon SUB1 depletion (Fig. 3C). We generated three independent viable U2OS SUB1 KO clones (ΔSUB1) (Fig. S3B). These clones had enhanced RPA32 pS33 at telomeres indicative of increased RS (Fig. 3D,E). ΔSUB1 cell lines also had increased telomeric BLM and higher numbers of APBs per nuclei (Fig. 3F and S3C,D). Conversely, RAD52 localization at telomeres, already high in normal U2OS cells, was not consistently altered in ΔSUB1 cells (Fig. S3E). As BLM dysregulation increases RS at ALT telomeres, we asked whether decreasing BLM levels might relieve some of the telomeric stress in ΔSUB1 cells (24, 25, 53–55). Indeed, using 2 independent siRNAs to deplete BLM resulted in a significant decrease of RPA32 phosphorylation at telomeres (Fig. 3G and S3F). We also noticed a slight but significant increase in telomere foci size in ΔSUB1 cells that could be attenuated by BLM KD, in agreement with the role of this helicase in telomere clustering during ALT (Fig. 3H) (56). Finally, because SUB1 functions at the interface between transcription and genome stability, we asked whether SUB1 influences TERRA levels at specific chromosome ends. We thus performed qRT-PCR on RNA extracted from U2OS and HeLa LT sgCtl and ΔSUB1 cells using chromosome-specific TERRA primer sets (57, 58).

**Figure 3.**
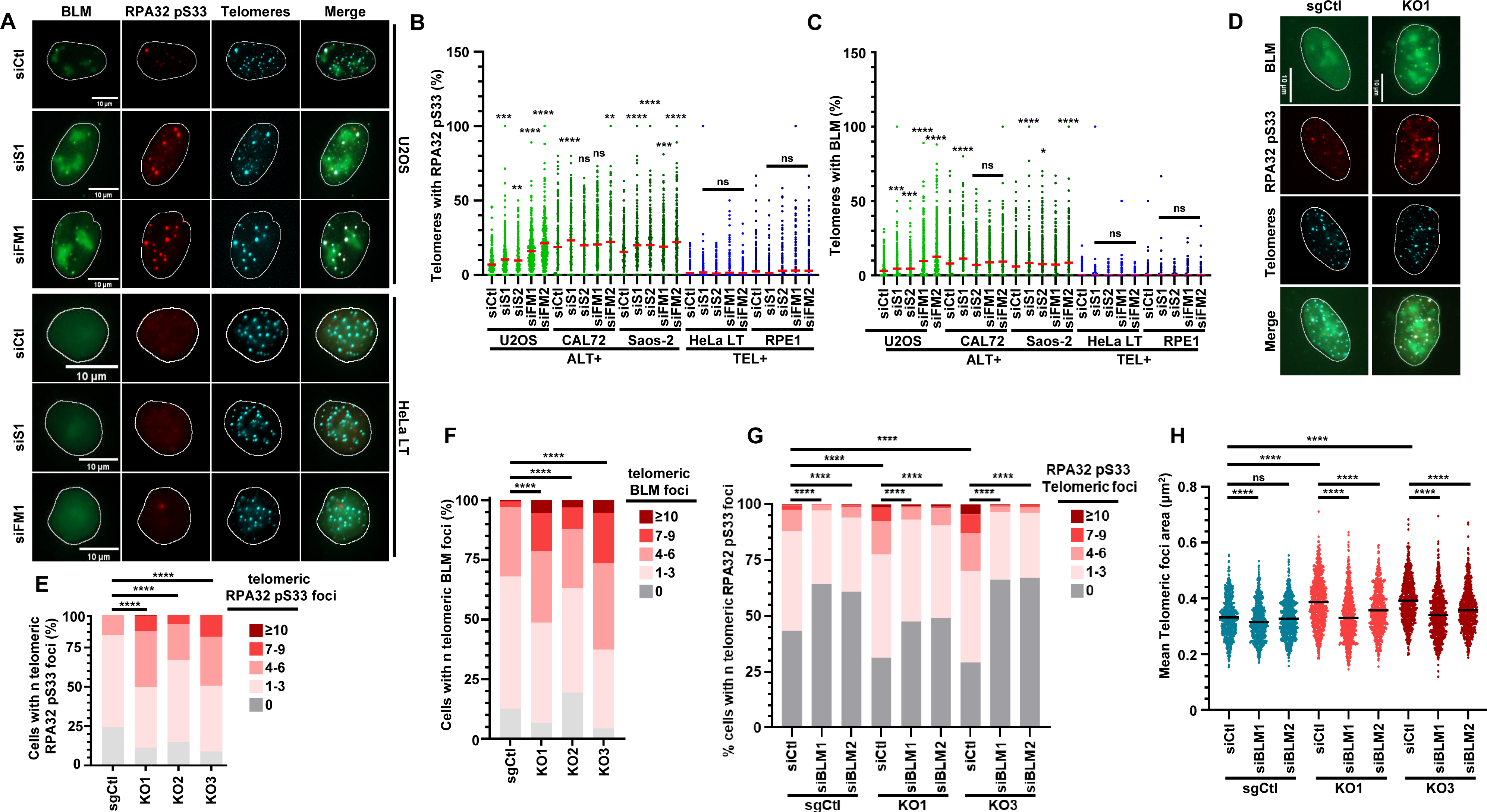
SUB1 alleviates RS at ALT telomeres. **(A-C)** A panel of ALT and TEL+ cell lines were transfected with the indicated siRNAs against SUB1 or FANCM and processed for RPA32pS33 and BLM IF along with telomeric FISH. **(A)** Representative IF-Tel-FISH in U2OS (ALT) and HeLa LT (TEL+). Quantification of RPA32pS33+ **(B)** or BLM+ **(C)** telomeres in ALT and TEL+ cell lines transfected with the indicated siRNAs was performed automatically using Cell Profiler. Graphs represent the merged data from three biological replicates. Statistical significance was established by one-way ANOVA and Holm-Šidák’s multiple comparison tests **(D-F)** sgCtl and KO SUB1 U2OS cells were processed for RPA32pS33 and BLM IF along with telomeric FISH. **(D)** Representative IF-FISH results for sgCtl and KO SUB1 cells. Histograms representing the percentage of cells with 0, 1-3, 4-6, 7-9 or ≥10 **(E)** telomeric RPA32 pS33 foci and **(F)** telomeric BLM foci. **(G,H)** sgCtl and KO SUB1 cells were transfected with the indicated siRNAs and 48 hrs later processed for Tel-FISH and IF. Graphs represent the merged data from three biological replicates. Statistical significance was established by Kruskal-Wallis and Dunn’s multiple comparison tests (P<0.05 *, P<0.01 **, P<0.001 *** and P<0.0001 ****).

We found that TERRA transcribed from multiple chromosomes was specifically enhanced in U2OS but not in HeLa LT ΔSUB1 cells suggesting a role for SUB1 in TERRA regulation (Fig. S3G). Altogether, these data suggest that SUB1 limits DNA RS and regulates several aspects of ALT telomere maintenance.

### SUB1 and FANCM Co-depletion is toxic in ALT Cells

As both FANCM and SUB1 alleviate RS at ALT telomeres and showed co-dependency in ALT contexts (Fig.1, S1), we tested whether they function epistatically or in parallel pathways to support ALT cell viability and telomere stability. We thus depleted FANCM and/or SUB1 in several TEL+ (HeLa LT, RPE1, HCT116) and ALT (U2OS, Saos-2, Cal72) cell lines and monitored colony formation over a 10-day period. Single depletion of SUB1 or FANCM using 2 independent siRNAs did not strongly affect the clonogenic potential of TEL+ cell lines compared to cells transfected with Ctl siRNA. In contrast, we observed a substantial decrease (∼50-70 %) in the number of colonies formed by ALT cell lines (Fig. 4A, S3A, S4A). These data are congruent with our initial Depmap analyses and previously published data on FANCM (Fig. 1A, B) (24–26). KD of SUB1 in the rhabdomyosarcoma ALT HS-729 cells also decreased their viability but did not impede the growth of non-ALT osteosarcoma cell lines 143b and HOS, further supporting the importance of SUB1 for ALT cell viability (Fig. S4B). Strikingly, co-depletion of SUB1 and FANCM in ALT cells led to an almost complete loss of viability (Fig. 4A). Depletion of FANCM in U2OS SUB1 KO cells also decreased cell viability suggesting that SUB1 KO and FANCM KD are particularly toxic for ALT cells (Fig. S4C,D). The decrease in viability in SUB1-depleted U2OS cells was readily rescued by re-expressing siRNA-resistant WT SUB1 but not the W89A or 3XMut (Fig. 4B, S4E), indicating that the ssDNA-binding function of SUB1 sustains ALT cell proliferation. The drop in ALT cell viability was accompanied by an increase in RPA32 pS33 foci and DNA damage levels at telomeres (γ-H2A.X) in SUB1/FANCM co-depleted ALT cells (Fig. 4C, S4F-H). Replication stress was also enhanced by SUB1 or FANCM KD in Hs729 ALT cells and RPA32pS33 intensity further increased upon co-depletion of these factors. Conversely, SUB1 and/or FANCM KD did not induce replication stress in non-ALT osteosarcoma cell lines HOS and 143b, confirming that SUB1 protects against replication stress specifically in ALT contexts (Fig. S4I, J). Furthermore, telomeric foci area and brightness increased in U2OS cells in response to FANCM depletion and this phenotype was exacerbated in SUB1/FANCM co-depleted cells (Fig. 4D, S4K, L) suggesting enhanced telomeric clustering in these cells. To probe the impact of SUB1 and/or FANCM on telomere structure, we performed in-gel telomere restriction fragment analysis of genomic DNA from U2OS and HeLa cells. SUB1 depletion in U2OS cells led to a decrease in G-rich ssDNA (Fig. 4E, F), possibly caused by shortening of the G-rich overhangs. A similar G-rich ssDNA decrease was seen upon FANCM abrogation as previously described (25, 26) and co-depletion of SUB1 and FANCM did not further enhance this phenotype (Fig. 4E,F, S4M). Strikingly, no such decrease was observed in HeLa Tel+ cells (Fig. 4E,F, S4M), and no alteration in overall telomere length could be detected upon SUB1 and/or FANCM depletion. As FANCM depletion was previously shown to drastically increase C-circle formation in ALT cells, we performed the C-circle assay using sgCtl or KO SUB1 U2OS or HeLa LT cell lines. KO of SUB1 in U2OS cells slightly but significantly increased C-circle levels (Fig. 4G, H), and depleting FANCM in U2OS KO SUB1 cells led to a further enhancement of this phenotype, suggesting that SUB1 may limit the production of extrachromosomal telomeric repeats in ALT cells when FANCM is abrogated (Fig. 4G, H). In line with these data, telomeric DNA synthesis was not detectably altered by SUB1 KO but FANCM depletion further enhanced EdU incorporation at telomeres in ΔSUB1 cells (Fig. S4N,O).

**Figure 4.**
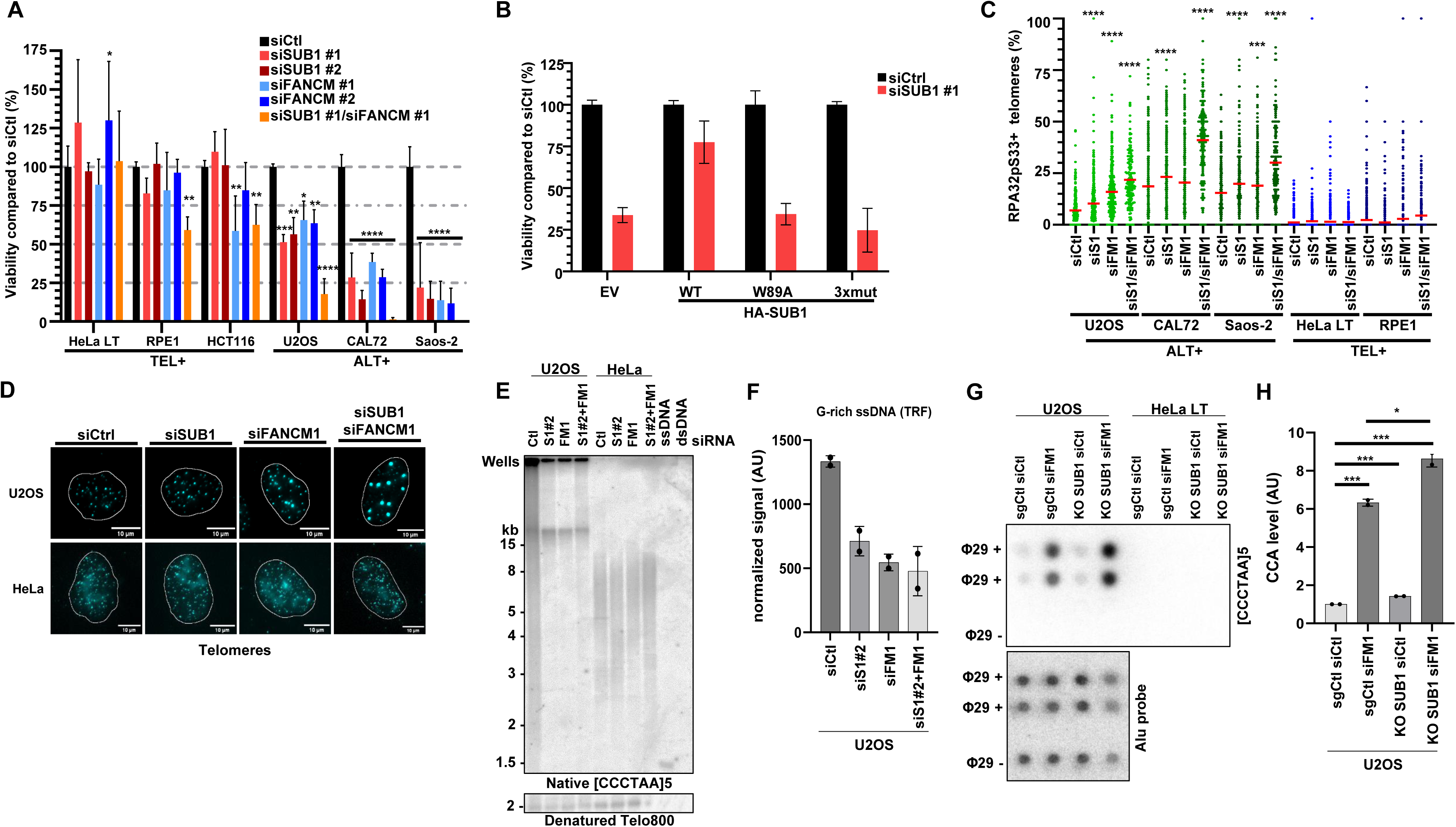
SUB1 is a vulnerability of ALT cells and its Co-depletion with FANCM is Highly Toxic. **(A)** U2OS, CAL72, Saos-2, HeLa LT or RPE1 cells were transfected with the indicated siRNAs. 500 cells per condition were seeded and colonies were counted after 10-14 days. Graph represents the merged data from two biological replicates each consisting of two technical replicates. **(B)** U2OS cells stably transduced with the indicated siRNA-resistant HA-SUB1 constructs were transfected with siRNAs and colony formation assays were performed 10-14 days post-transfection. Graph represents the merged data from two biological replicates each consisting of two technical replicates. **(C)** U2OS, CAL72, Saos-2, HeLa LT or RPE1 were transfected with the indicated siRNAs and processed for immunofluorescence and telomere FISH. Telomeric RPA32pS33 foci were automatically counted using CellProfiler. The data presented for siCtl, siS1 and siFM1 are the same as in Fig. 3B as SUB1/FANCM co-depletion experiments were performed simultaneously. Experiments were performed in biological duplicates and a representative replicate is shown here. Statistical significance was established by one-way ANOVA and Holm- Šidák’s multiple comparison tests **(D)** Representative images of telomeric foci in U2OS and HeLa cells. **(E,F)** G-rich ssDNA TRF analysis of ALT (U2OS) and telomerase+ (HeLa) siRNA-transfected cells. Genomic DNA was extracted 48 hours after siRNA transfection, digested with a restriction enzyme mix, and underwent native in-gel hybridization with radiolabeled [CCCTAA]5 oligonucleotides targeting the G-rich overhang. A denaturing Southern blot with a radiolabeled telomeric probe (Telo800) was used for normalization. ssDNA and dsDNA controls of TRF enzyme-digested from a plasmid indicate that the probe hybridized only to ssDNA. Experiments were performed in biological replicates **(G,H)** C-circle assay (CCA) analysis of genomic DNA from the indicated sgCtl or SUB1 KO U2OS or HeLa LT cell lines transfected with control or FANCM-targeting siRNAs. Reactions were performed with or without Phi29 polymerase (Φ29+/-). CCA products were dot-blotted and hybridized with a radiolabeled [CCCTAA]5 oligonucleotide probe. C-circle values were quantified relative to siCtl. Biological replicates were performed. Statistical significance was established by one-way ANOVA and Holm-Šidák’s multiple comparison tests (p-values: P<0.05 *, P<0.01 **, P<0.001 *** and P<0.0001 ****).

**Figure 5.**
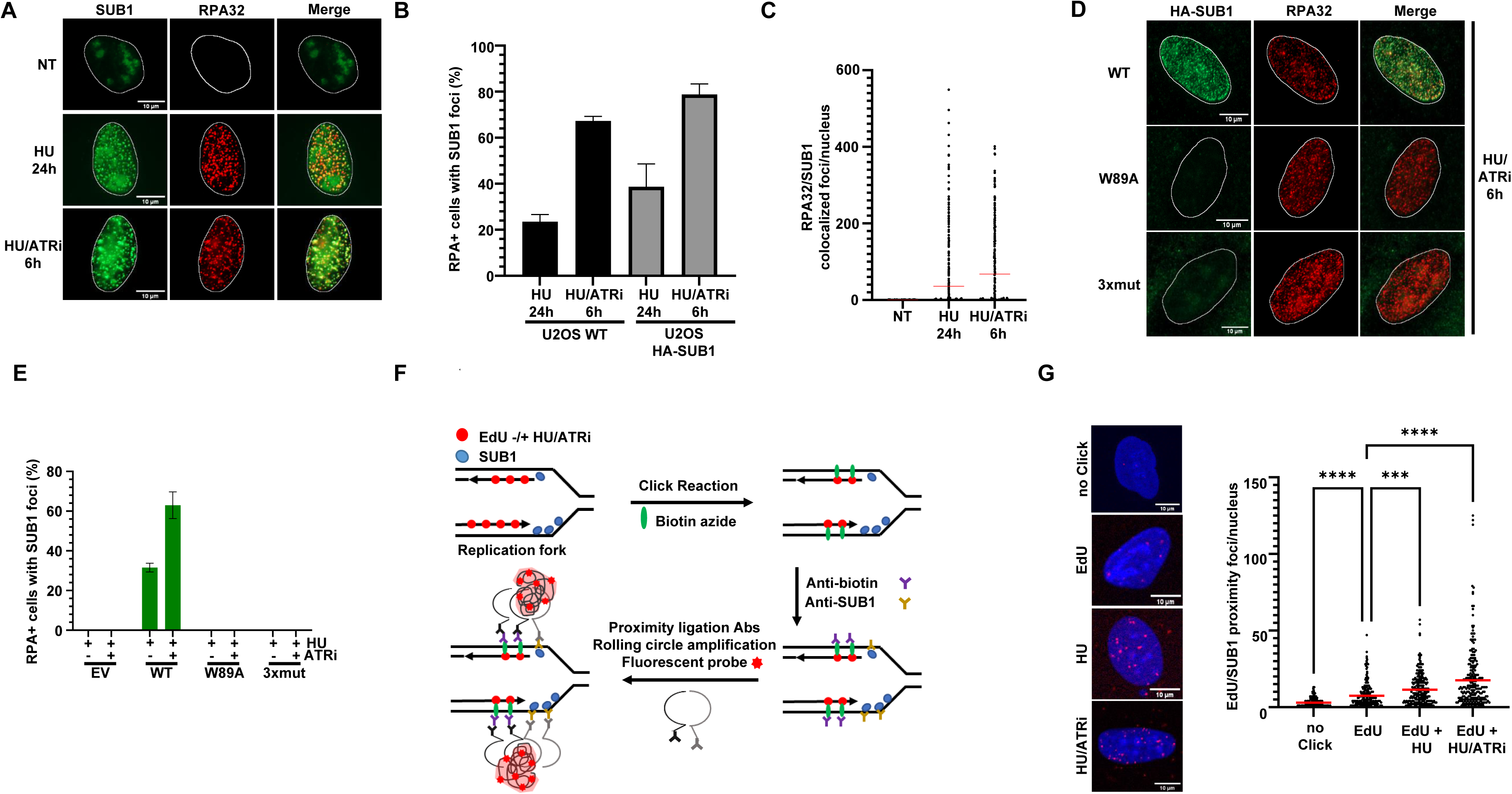
The single-stranded DNA binding protein SUB1 associates with stalled and collapsed DNA replication forks. **(A-C)** U2OS WT or HA-SUB1 cells were treated with 2mM HU or HU combined with 1 uM ATRi for the indicated times. After fixation, IF against SUB1/HA and RPA32 was performed. RPA32- and SUB1-foci were automatically counted using CellProfiler. **(A)** Representative images of WT U2OS cells stained for endogenous SUB1 and RPA32 IF. Nuclei were outlined with the “Outliner” ImageJ plugin. **(B)** Histogram representing the fraction of HU- and HU/ATRi-treated cells containing SUB1 foci. Cells with >30 RPA and SUB1 foci were considered positive. **(C)** Graphical representation of the number of foci contained in each nucleus. Graphs represent the merged data from two biological replicates **(D)** SUB1 mutants were stably transduced into U2OS cells that were then treated with HU and ATRi. Representative immunofluorescence images are shown. **(E)** Graphs represent the merged data from two biological replicates **(F)** Schematic representation of *in situ* analysis of protein interactions at DNA replication forks (SIRF). **(G)** Cells were exposed for 30 minutes to EdU (100 uM) to label ongoing replication forks. Cells were then treated or not with HU or a combination of HU and ATRi to induce fork stalling and collapse respectively. Cells were processed for SIRF as outlined and EdU-SUB1 proximity foci were automatically acquired using CellProfiler. Statistical significance was established by one-way ANOVA and Šidák’s multiple comparison tests. Experiments were performed in biological triplicates and a representative replicate is shown here (P<0.001 ***, P<0.0001 ****).

### SUB1/PC4 Localizes to Stalled and Collapsed DNA Replication Forks

Apart from its role at ALT telomeres which we describe here, SUB1 was proposed to play a more general role in the mitigation of RS at stalled forks (40, 41). To test this model, we confirmed, using HA-tagged SUB1 or using antibodies recognizing endogenous SUB1, that inducing RS by depleting nucleotide pools with hydroxyurea (HU) and inhibiting ATR (ATRi) led to the formation of bright SUB1 foci (Fig. 5A, B and Fig S5A-E). Consistent with previous data (40), these foci significantly colocalized with RPA (Fig. 5C). Recruitment was dependent on an intact SUB1 ssDNA-binding domain as in cells expressing both the W89A and the 3XMut constructs, SUB1 foci could not be detected anymore (Fig. 5D, E). Interestingly, upon ATRi treatment, RPA foci formation occurred prior to SUB1 foci appearance, suggesting that RPA may precede SUB1 loading on ssDNA (Fig. S5F). Finally, proximity-ligation assays in EdU-treated cells showed that SUB1 associates with ongoing replication forks in naïve cells and this was enhanced by HU and even more so by ATRi/HU co-treatment indicative of an association with stalled forks (Fig. 5F, G).

### SUB1 Protects against RS and Replication Catastrophe in ALT Cells

The association of SUB1 with ongoing and stalled forks and the enhanced telomeric RS that occurs in SUB1 and FANCM co-depleted cells prompted us to test whether SUB1 may also help ALT cells cope with other sources of RS. Indeed, SUB1 depletion sensitized U2OS cells to HU, ATRi and PARPi treatments (Fig. 6A-C, S6A). Similarly, SUB1 KO cells were also more sensitive to ATR and PARP inhibition (Fig. S6B,C). In contrast, depleting SUB1 in HeLa LT cells did not significantly alter their sensitivity to these agents, suggesting that ALT cells may be more reliant on SUB1 to overcome RS (Fig. S6A, D-F).

**Figure 6.**
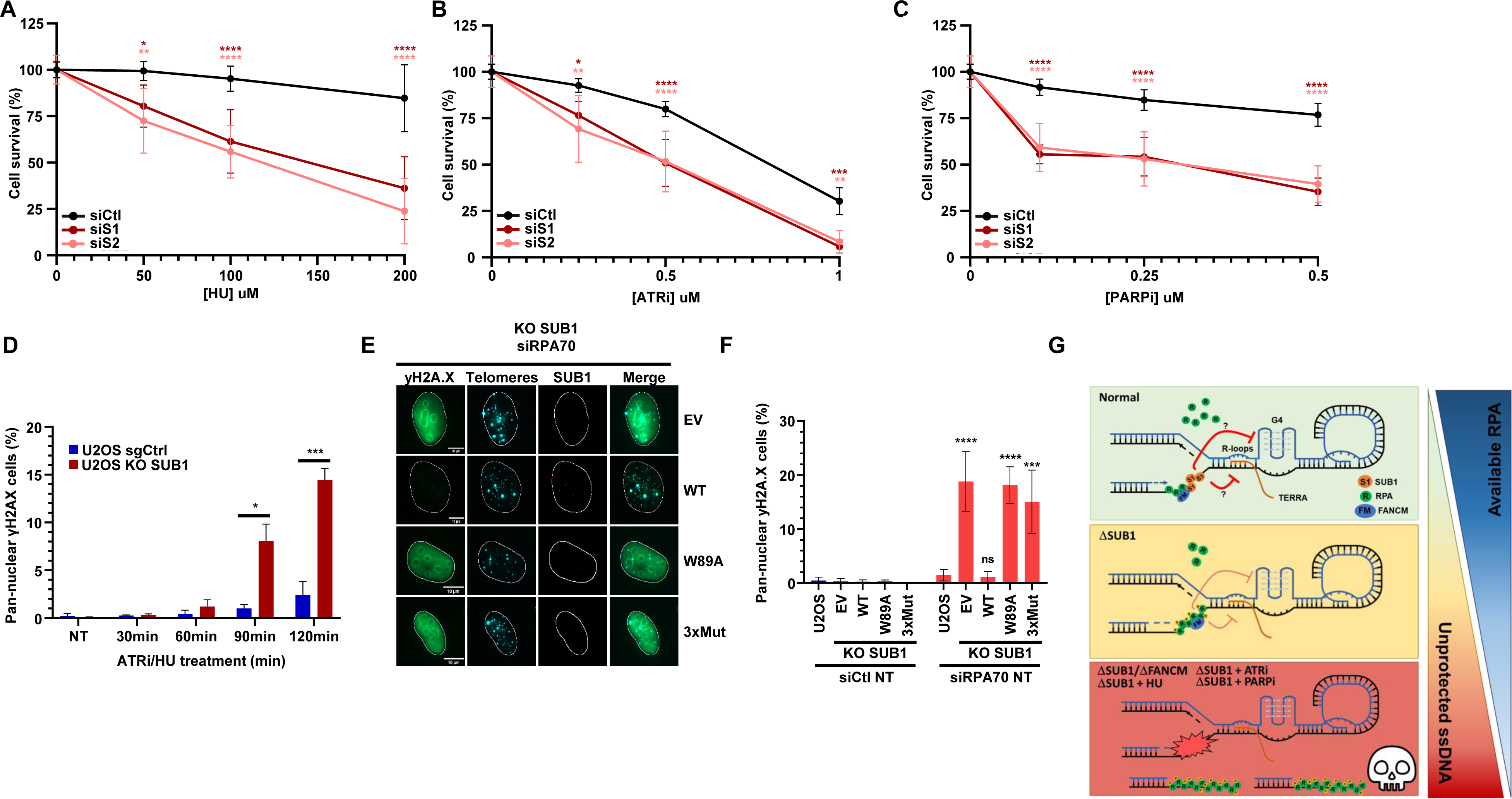
SUB1 Protects Against RS and Replication Catastrophe in ALT cells. **(A,B,C)** U2OS cells were transfected with the indicated siRNAs and incubated with HU, ATRi or PARPi at the indicated concentrations for 4 days. Cells were then released into fresh media and allowed to form colonies over 10 days prior to fixation and viability assessment. Graphs represent cell viability compared to non-treated cells. Statistical significance was established by ordinary two-way ANOVA and Tukey’s multiple comparison tests **(A)** Graphs represent the merged data from four biological replicates each consisting of two technical replicates. **(B-C)** Graphs represent the merged data from three biological replicates each consisting of two technical replicates. **(D)** U2OS sgCtl and KO SUB1 cells were exposed to ATRi and HU for the indicated times and processed for γ-H2A.X immunofluorescence. γ-H2A.X nuclear intensity were automatically counted using Cell Profiler. Cells with nuclear γ-H2A.X ≥250 were considered pan-nuclear. Experiments were performed in biological duplicates and a representative replicate is shown here. **(E)** Representative images and **(F)** quantification of pan-nuclear γ-H2A.X cells. WT and KO SUB1 cells complemented with the indicated HA-SUB1 constructs were transfected with the indicated siRNAs and 48 hrs late processed for γ-H2A.X and SUB1 IF along with telomere FISH. Graph represents the merged data from three biological replicates. Statistical significance was established by ordinary two-way ANOVA and Tukey’s multiple comparison tests **(G)** A model for the roles of SUB1 in the resilience of ALT cells towards replication stress. In ALT cells, SUB1 binds ssDNA at stressed telomeres and ongoing replication forks and collaborates with RPA, FANCM and other factors to alleviate RS at ALT telomeres and elsewhere in the genome. Due to its roles at the interface between transcription and replication, its affinity for G4-forming DNA and its potential role in limiting TERRA levels, SUB1 might help mitigate R-loop and G4 formation at ALT telomeres. SUB1 depletion enhances RS at telomeres, making ALT cells more dependent on RPA and prone to undergo replication catastrophe. Co-depleting SUB1 and FANCM or combining SUB1 depletion with RS-inducing drugs is highly toxic to ALT cells. (P<0.05 *, P<0.01 **, P<0.001 *** and P<0,0001****).

Prolonged ATR inhibition depletes available nuclear RPA pools, eventually leading to exposed ssDNA and rampant fork breakage, a phenomenon known as replication catastrophe (59–61). To determine whether SUB1 may help cells avoid fork breakage during prolonged RS, we monitored the induction of pan-nuclear γ-H2A.X, a hallmark of replication catastrophe in cells treated with ATRi and HU (60). KO SUB1 cells accumulated pan-nuclear γ-H2A.X at a higher magnitude and faster than sgCtl U2OS cells (Fig. 6D), suggesting that SUB1 may make cells more resilient towards RPA exhaustion during acute RS. The relationship between RPA and SUB1 during RS also suggests a model in which SUB1 may assist RPA to coat and protect ssDNA, thereby preventing fork collapse and DSB induction (Fig. 2 and S5F, and (40)). To formally test this idea, we depleted RPA for 48 hrs in sgCtl U2OS cells, which led to a very mild and non-significant increase in pan-nuclear γ-H2A.X cells (Fig 6E,F). In sharp contrast, RPA depletion led to ∼20 % of KO SUB1 cells containing bright pan-nuclear γ-H2A.X staining (Fig. 6E,F), indicating enhanced sensitivity to RPA exhaustion. These phenotypes along with ATR activation monitored by CHK1-pS345 were rescued by WT SUB1 but not ssDNA-binding mutants (Fig. 6E, F, S6G), suggesting that SUB1 may interact with ssDNA to limit replication stress and prevent catastrophic fork cleavage when RPA levels become limiting. In further support of this model, re-expression of WT SUB1 but not the W89A mutant also significantly suppressed the accumulation of RPA at telomeres in ATRi/HU-treated SUB1 KO U2OS cells (Fig. S6H).

## Discussion

Here, we have leveraged recent CRISPR fitness screening efforts hosted on the Cancer Dependency Map to identify genetic vulnerabilities of ALT cancer cells (62). The approach identified several expected genes previously shown to be involved in ALT cell viability and/or telomere maintenance, including the FANCM helicase complex (24–26), the SMARCAL1 DNA translocase (23, 27) and the FANCA, FANCF and FANCG components of the Fanconi anemia core ubiquitin ligase complex (26, 36). Additionally, we identified the RAP1/TERF2IP shelterin protein as a putative ALT vulnerability. Notably, RAP1 is known to limit homologous recombination engagement at telomeres (63, 64). Whether RAP1 is relevant to ALT cell viability will require further investigation. Our analyses generally agree with and build upon those from a recent publication that identified a similar set of ALT-specific vulnerabilities (35). A commonality of most of the dependencies identified here is that they alleviate telomeric RS and limit recombinogenic activity at telomeres. We present robust evidence that the ssDNA-binding factor SUB1/PC4 is a novel factor that mitigates telomeric RS to support ALT cell proliferation.

Extending upon prior findings, we show that SUB1 is present at ongoing and stalled replication forks and forms telomeric foci almost exclusively in S-phase cells (Figs. 2F,5, S5 and (40, 41)). These data are further corroborated by recent unbiased chromatin capture and iPOND proteomics which found SUB1 at ongoing or stalled replication forks, respectively (65, 66). In cells challenged with ATRi, SUB1 is also attracted to ALT telomeres via its ssDNA-binding domain, indicating that SUB1 interacts with ssDNA both at stalled forks and damaged telomeres (Figs. 2, S2, 5 and S5). Interestingly, RPA depletion greatly enhanced SUB1 recruitment to telomeres and at stalled forks. Conversely, re-expression of WT but not W89A mutant SUB1 in SUB1 KO U2OS cells decreased RPA accumulation at telomeres upon ATRi/HU co-treatment (Fig. S6H). These experiments suggest that SUB1 and RPA may bind the same substrates, namely ssDNA during RS, and that this competition may be more acute in ALT contexts, increasing the need for SUB1 in this situation. In agreement with these data, a tug-of-war between these 2 avid ssDNA-binders occurs during transcription in yeast and in *in vitro* SV40 virus replication assays (67, 68). Antagonism between RPA and SUB1 was also observed by the Helleday lab who proposed that SUB1 present at transcription bubbles may relocate onto RPA-coated ssDNA during replication-transcription collisions to promote genome stability (40). Our data shows that SUB1 is particularly important to resolve RS in the context of ALT telomeres, which contain high levels of ssDNA particularly prone to forming structures that impede replication and thus may require a lot of RPA (Figs. 3, 4, S4 and 6G). Indeed, we demonstrate that RPA is recruited prior to SUB1 at stalled forks and that SUB1 helps avoid replication catastrophe by binding ssDNA when RPA becomes limiting (Figs. 6E, F, S5F and S6G). How SUB1 influences other ssDNA-binding factors such as POT1/TPP1 or the CST complex at telomeres will be interesting areas for future research. Altogether, our work suggests a model in which SUB1 collaborates with RPA to minimize RS and protect ssDNA at telomeres and genome-wide (Figs. 6, S5F and S6).

Telomeres are predisposed to guanine oxidation and readily form G4s and TERRA RNA:DNA hybrids; potent obstacles to DNA polymerases and key regulators of break-induced replication in ALT cells (69–74). There is evidence that SUB1 could be involved in the modulation of each of these structures, potentially explaining its role in RS mitigation at ALT telomeres. First, Sub1 inactivation in yeast sensitizes cells to reactive oxygen species and this function can be complemented by human SUB1. Purified Sub1 also binds to and protects DNA against oxidative damage *in vitro*, raising the possibility that it might directly counteract guanine oxidation at telomeres as well (75). Secondly, yeast and human SUB1 bind G4-forming DNA sequences including human telomeric repeats with affinities similar to or higher than for ssDNA. Mutational analyses also revealed that key residues of the ssDNA-binding surface (F77 or W89) are important for association with G4 and ssDNA substrates, suggesting that dimeric SUB1 may interact simultaneously with ssDNA and G4s (76, 77). In contrast to RPA, SUB1 is unable to unfold pre-formed G4s on its own but does slow down the folding rate of these structures *in vitro* (77, 78). In *Saccharomyces cerevisiae,* Sub1 also represses transcription-associated G4-mediated recombination and this function can be taken over by the ssDNA-binding domain of human SUB1 in complementation assays (44). It is thus possible that SUB1 could directly modulate the kinetics of G4 formation and/or recruit additional factors to help dissolve these structures, thereby relieving telomeric RS. Finally, SUB1 functions at the interface between replication and transcription (37). ALT telomeres have abundant levels of TERRA RNA:DNA hybrids that need to be regulated for optimal telomere synthesis. Intriguingly, we found that TERRA originating from several chromosome ends was more abundant in U2OS ΔSUB1 cells compared to the sgCtl but that SUB1 inactivation in TEL+ HeLa cells did not affect TERRA levels (Fig. S3G). This suggests that SUB1 may regulate the transcription and/or stability of specific TERRA RNA species in ALT cells. What the role(s) of SUB1 are in this process and whether the observed increase in TERRA is a direct or indirect consequence of SUB1 inactivation will be important to determine. In addition to regulating TERRA levels, SUB1 could conceivably interact with extruded ssDNA in RNA:DNA hybrids and modulate their formation at ALT telomeres (58, 79, 80). It has been previously shown that FANCM limits RS at ALT telomeres at least in part via the modulation of BLM activity within APBs (24, 25). Similar to FANCM depletion, SUB1 KD or KO led to an accumulation of BLM at ALT telomeres and depletion of this helicase alleviated telomeric RS and clustering in SUB1 KO cells (Figs. 3C, F, G, H and S3F). Moreover, BLM depletion specifically enhanced the proliferation of SUB1 KO compared to sgCtl U2OS cells strongly suggesting that ALT telomere RS induced by SUB1 depletion arises at least partly via BLM deregulation (Fig. S7). More investigations will be required to determine exactly which source(s) of telomeric RS is/are regulated by SUB1 and determine their respective contributions to ALT cell death when SUB1 is impaired.

Most importantly, our work shows that SUB1 depletion is specifically deleterious to ALT cells. We also establish a toxic genetic relationship with the FANCM helicase, with co-depleted cells having greatly exacerbated markers of telomeric RS and damage and accumulating large clusters of telomeres indicative of deregulated maintenance pathways (Figs. 4, and S4). SUB1 depletion also synergizes with RS-inducing drugs and accelerates replication catastrophe when RPA becomes limiting, a phenomenon that appears to be particularly effective in ALT backgrounds (Figs. 6 and S6). The high vulnerability of ALT cells towards RS likely stems from a combination of factors including their tendency to form telomeric secondary structures and their use of break-induced replication pathways to elongate telomeres (4, 14, 30, 81). ALT cells also generally have higher levels of chromosomal and extrachromosomal telomeric ssDNA than TEL+ cells (18, 82–85). Such abundant single-stranded species could act as sinks for ssDNA-binding factors, decreasing the threshold at which RPA becomes limiting and replication catastrophe is induced. In such contexts, ssDNA-binders such as SUB1 and other factors that limit RS at telomeres (FANCM, SMARCAL1, ATR…) may become necessary to sustain viability. Our data indeed suggest that SUB1 can partially replace FANCM in cases where the latter is absent: SUB1 loss does not lead to increased extrachromosomal C-circles, yet its absence is felt if FANCM is limiting (Fig. 4G, H). In the same vein, telomeric DNA synthesis is not substantially increased in SUB1 KO cells but we observed a synergistic impact of FANCM depletion on telomeric EdU incorporation specifically in ΔSUB1 cells (Fig. S4N, O). Therefore, while FANCM normally is sufficient to limit C-circle generation, SUB1 can partly perform this role in its absence. Remarkably though, the level of telomeric ssDNA overhangs is similarly reduced by an absence of either gene, suggesting that on chromosome ends, the two proteins play entirely redundant roles (Fig. 4E, F). We propose that SUB1 constitutes an alternative reservoir of ssDNA protection that buffers RS in a variety of settings, including at ALT telomeres. Under conditions of mild RS at telomeres, FANCM and RPA are able to mitigate some of the stress surge that occurs upon SUB1 depletion. However, when RS levels are increased exogenously above a certain threshold either by depleting ALT regulators or by exposure to genotoxic drugs, SUB1’s role becomes essential to sustain telomere and genome stability to support cell viability (Fig. 6G). Altogether, we have characterized SUB1 as a novel vulnerability of ALT cells with key roles in telomere stability and provide several new synthetic sick relationships that may potentially be therapeutically exploited to improve the treatment outcomes of a group of rare but deadly cancers.

## Material and Methods

### Cell Lines and Culture Conditions

HCT116, U2OS, HeLa, HeLa LT, Saos-2, CAL72, HEK293T, Hs729, HOS and RPE1 hTERT cells were obtained from ATCC. 143b cells were obtained from Riken. Cell lines were grown in Dulbecco’s modified Eagle’s Medium (DMEM) supplemented with 1% streptomycin/penicillin antibiotics (Wisent) and 10% fetal bovine serum (Gibco). Cells were grown at 37°C in a 5% CO2 humidified atmosphere. Cells in culture were regularly tested to ensure the absence of mycoplasma contamination. For treatments, hydroxyurea (HU) (Bioshop), AZD6738 (ATRi) (Adooq Bioscience), Olaparib (PARPi) (Selleck Chemicals) were used as indicated in the corresponding figure legends. For cells synchronization, cells were treated with 2mM thymidine for 20-24hrs, released for 9hrs and retreated with 2mM thymidine for 18-20hrs. Cells were then released for 0hrs (G1 phase), 5hrs (S phase) or 9hrs (G2 phase).

### Antibodies, Reagents and Oligonucleotides

Key reagents used in this study are presented in Table S9.

### CRISPR Fitness Screens Analyses

Experimentally validated ALT cancer cell lines were used as a representative ALT cell group SaOS2, U2OS, CAL72, CAL78, HuO9, SKNFI G292CLONEA141B1. DEPMAP (21Q1) CERES scores for all cell lines were downloaded and mean subtracted scores (differential dependencies) were computed for all cell lines in the database (CERES score for gene X in cell line Y – average CERES score for gene X across the full panel of cell lines). The average differential dependencies for the ALT cell lines group (n = 7) were then calculated for all genes and unpaired t-tests were performed to identify statistically significant differential dependencies in the ALT cell lines group (Table S1). These analyses were also performed using raw CERES scores for ALT vs non-ALT cells (Fig. S1A) and extended to a larger panel of ALT cell lines (n = 24) on which CRISPR fitness screens had been performed (Table S7) (43). Differential dependencies were also identified using the top 5 % (Z-score <-1.64) FANCM-(n = 40) and FANCM complex-dependent (n = 48 (mean differential dependency scores for FANCM-MHF1/2-FAAP24)) (Tables S3,4). For co-dependency analyses, Pearson correlations were performed between SUB1 and all other genes using raw CERES (Depmap 21Q1) or CRISPR (Depmap 23Q2) scores.

### Plasmids and Mutations

SUB1 WT, W89A and 3XMut resistant to siSUB1 #1 siRNA cDNA sequences were synthesized by Eurofins. PCR were performed using AttB1-SUB1 and AttB2-SUB1-stop oligonucleotides and subsequently transferred into pDONR221 entry and pEF1a-3xHA destination vectors using Gateway cloning (Thermo-Fisher). pLenti-U6-sgRNA-SFFV-Cas9-2A-Puro containing 3 different sgRNAs against SUB1 were obtained from ABM goods.

### Generation of SUB1 KO cells and cell lines stably expressing HA-SUB1 constructs

Lentiviruses were produced by standard methods. Briefly, HEK293T cells at 80 % confluence were co-transfected with pHAGE EF1α-3XHA lentiviral vector, VSV-G envelope expressing plasmid pMD2.G (Addgene # 12259) and lentiviral packaging plasmid psPAX2 (Addgene # 12260) by the polyethyleneimine method. Supernatants containing viruses were collected and filtered (0.45-µm pores) 48 h post-transfection. Infections of U2OS/HeLa LT cells were performed in the presence of polybrene (hexadimethrine bromide, Sigma). Stable cell lines were produced using puromycin selection. KO was confirmed by immunoblotting and Sanger sequencing of PCR products from genomic DNA and CRISP-ID allelic deconvolution (86).

### siRNA Knockdowns

Reverse transfection of siRNAs were performed using Lipofectamine RNAiMAX according to manufacturer’s recommendations (ThermoFisher/Life Technologies). Briefly, siRNA were diluted in OptiMEM and incubated for 3 minutes. Lipofectamine RNAiMAX was then added to diluted siRNA and the reaction was incubated at RT for at least 15 minutes before transfection.

### Immunofluorescence and Telomere FISH

Cells were seeded onto coverslips and treated as described in the corresponding figure legends. Cells were washed with PBS1X and pre-permeabilized with CSK buffer (10mM Hepes-KOH [pH 7.4], 300mM sucrose, 50mM NaCl, 3 mM MgCl2, 1mM EGTA, 0.5% Triton X-100) for 5 minutes at 4°C (except for expression level experiments which were performed without pre-permeabilization). Cells were then rinsed with PBS1X and fixed/permeabilized with PTEMF buffer (20 mM PIPES pH 6.8, 10 mM EGTA, 0.2% Triton X-100, 1 mM MgCl2 and 4% formaldehyde) for 20 minutes at room temperature (RT). Cells were rinsed with PBS1X and incubated in blocking buffer (0.05% tween, 3% BSA in PBS1X) for 30 minutes at RT. Coverslips were incubated overnight at 4°C in a humidified chamber with the primary antibodies diluted in blocking buffer. Coverslips were washed 3 times, 5 minutes with PBS1X-Tween 0.05 % and agitation and incubated 1h 30 at 37 °C in a humidified chamber with fluorescent secondary antibodies (AlexaFluor 488 and AlexaFluor 647). For experiment without FISH hybridization, samples were washed 3 times, 5 minutes with PBS1X-Tween 0.05% and agitation, incubated in PBS1X containing DAPI (1ug/mL) for 10 minutes with agitation, rinsed with PBS1x and mounted onto slides using Prolong Diamond Antifade mounting media (Thermo Fisher/Life Technologies). For FISH hybridization, samples already hybridized with primary and secondary antibodies were fixed with PFA 4% for 10 minutes at RT. Samples were then rinsed with PBS1X and incubated in SSC2X (300mM NaCl, 30mM sodium citrate, pH7) containing 200ug/mL RNAseA (Monarch) for 30 minutes at 37°C in a humidified chamber. Samples were rinsed in PBS1X and subsequently dehydrated in ethanol 70%, 85% and 100% for 2 minutes each at RT. Coverslips were then dried at 37°C for 15 minutes. Samples were incubated in pre-heated FISH-hybridization buffer (410µM Na2HPO4, 125µM MgCl2, 450µM citric acid, 10mM Tris pH7.5, 2.5 mg/mL blocking reagent (Roche) in 70% formamide v/v (Sigma, Fisher)) at 85°C for 5 minutes and put in a slightly humidified chamber at 37°C overnight for hybridization. Samples were then washed 3 times, 5 minutes with SSC2X:Formamide 1:1 solution and 3 times with SSC2X at 37°C with agitation. Coverslips were then incubated in SSC2X containing DAPI (1ug/ml) for 10 minutes with agitation and mounted on slides using Prolong Diamond Antifade mounting media (Thermo Fisher/Life Technologies). Images were collected using a fluorescence Zeiss Axio Observer Z1 microscope with Zeiss Axiocam 506 mono camera. Zeiss Zen 2.0 software was used to capture images. Images were processed with Fiji and analysed automatically with CellProfiler (87) for signal and foci quantification.

### Image processing and analysis using CellProfiler

Illumination correction was initially applied to the images using CellProfiler. For each image, a function representing background as well as the uneven illumination was calculated and subsequently subtracted from the image. To detect foci, images were first processed to enhance speckles using a top hat filter with a diameter equal to the largest visible foci. Segmentation of the processed images was performed using a manual threshold, and objects touching each other were split by detecting intensity peaks. Each intensity peak was considered a single object. Intensity measurements were done on the images with corrected illumination. Colocalized foci were determined using a method established by the CellProfiler developers (https://cellprofiler.org/examples). For each channel where foci were identified, each detected object was shrunk to a single pixel and then expanded by two pixels. Objects still overlapping were considered colocalized.

### Telomere DNA synthesis assay

Cells were seeded onto coverslips and treated with 10µM EdU for 20-30 minutes 48h after appropriate knockdown. Cells were then washed with PBS1X and pre-permeabilized with CSK buffer (10mM Hepes-KOH [pH 7.4], 300mM sucrose, 50mM NaCl, 3 mM MgCl2, 1mM EGTA, 0.5% Triton X-100) for 5 minutes at 4°C. After CSK incubation cells were rinsed with PBS1X and fixed/permeabilized with PTEMF buffer (20 mM PIPES pH 6.8, 10 mM EGTA, 0.2% Triton X-100, 1 mM MgCl2 and 4% formaldehyde) for 20 minutes at room temperature (RT). After rinsing cells with PBS1X, a Click reaction was performed by incubating coverslips on Fluor-azide cocktail (100mM Tris HCl pH 8.5, 1mM CuSO4, 20µM Ethereon-Red 645 azide and 100mM ascorbic acid) for 1 hour at RT. Cells were then rinsed with PBS1X and IF protocol was performed from blocking solution onwards as described earlier to combine with TRF2 detection.

### *In situ* protein interaction with nascent forks

Experiments were performed as previously described (88). Briefly, cells seeded on coverslips were pulsed with 100mM EdU for 30 minutes and treated or not with HU 5mM or 2mM HU/1uM ATRi for 4 hrs. After treatment, cells were washed with PBS1X and incubated in pre-extraction buffer (20 mM NaCl, 3 mM MgCl2, 300 mM sucrose, 10 mM PIPES, 0.5% Triton X-100) for 10 minutes on ice. Cells were then washed with PBS1X and fixed in PFA 3%, 2% sucrose for 15 minutes at RT. After fixation, cells were washed with PBS1X and permeabilized in 0.25% triton X-100 for 15 minutes on ice. Cells were washed in PBS1X and incubated in 10% BSA solution overnight at RT. Cells were then washed with PBS1X and coverslips were incubated on Click-it solution (20 μM biotin azide, 10 mM sodium ascorbate, 2 mM CuSO4 in PBS1X) for 1h at RT. Samples were washed in PBS1X and blocked in 10% BSA solution for 30 minutes. Coverslips were then incubated on primary antibodies (α-Biotin and α-SUB1) diluted in PBS1X 10% goat serum, 0.05% tween overnight at 4°C in an humidified chamber. PLA protocol was then performed as per manufacturer’s instructions (Sigma, Duolink PLA kit). Images were collected using a fluorescence Zeiss Axio Observer Z1 microscope with Zeiss Axiocam 506 mono camera. Zeiss Zen 2.0 software was used to capture images. Images were processed with Fiji and analysed with CellProfiler for proximity foci quantification.

### Colony formation assays

Cells were seeded onto siRNA-lipofectamine complexes and 48 hrs later (or 24h for RS sensitivity experiments), cells were harvested by trypsinization for immunoblotting and reseeding onto 6-well plates at 500 cells/well (1000 for RS sensitivity experiments) with two wells technical replicates per condition. For RS experiments, cells were treated for 4 days with drugs 24h after reseeding. Colonies were left to grow for 10-14 days and fixed in ethanol 70% at 4°C for at least 20 minutes. After fixation, colonies were stained with crystal violet for 15 minutes on a shaker at room temperature. Viable colonies were then counted by hand for each condition.

### Native in-gel hybridizations and Southern blots

Cell lysates were treated with 20 µg/mL RNase A and 100 µg/ml Proteinase K, DNA was isolated by phenol:chloroform:isoamyl (25:24:1) extraction and precipitated with 100% ethanol + 300 mM NaOAc. Resuspended DNA was digested with a restriction enzyme mix of *Hinf*I, *Rsa*I, and MboI (NEB) and quantified by fluorometry. For TRF analysis, 7.5 μg of digested DNA was separated on 0.7% 1X TBE agarose gels and vacuum dried for approximately 30 min with a Bio-rad gel dryer (model 583). Gels were sealed in a plastic bag and hybridized at 37 °C overnight with a 5′-(CCCTAA)5−3′ telomeric oligonucleotide probe 5′-end-labeled with T4 polynucleotide kinase (NEB M0201S) and [γ-^32^P]ATP (PerkinElmer BLU502A). Gels were washed 3-4 X 30 min in 0.25X SSC at RT and exposed on a phosphor cassette. The native gels were then subjected to denaturing Southern blot to visualize the total telomeric DNA and internal telomeric repeat bands. The Southern blot was performed as follows: 10 minutes in 0.25M HCl, 45 minutes in Southern denaturing solution (1.5M NaCl, 0.5M NaOH), 5 minutes 0.4M NaOH, and transferred to a Hybond-XL nylon membrane (Cytiva). The membrane was then hybridized at 65°C overnight with a double-stranded 800 bp telomeric repeat probe (Telo800) that was radioactively labeled with Klenow (NEB M0210S) and [α-^32^P]dCTP (PerkinElmer BLU513H). The membrane was washed for 20 min in 2X SSC at RT and exposed on a phosphor cassette. Radioactive signal imaging was performed with a Typhoon FLA 9000 imager (GE Healthcare) and quantified using ImageJ software.

### C-Circle Assays

100 ng of extracted and digested genomic DNA was mixed with 7.5 U phi29 DNA polymerase (NEB M0269S), and 1 mM each of dATP, dTTP, and dGTP to a total volume of 20 µl. The reaction was incubated at 30 °C for 4 h or 6 h and heat-inactivated at 65 °C for 20 min. Amplification products were diluted to 50 µl in nucleotide-free water, dot-blotted (Bio-rad 1706545) onto Hybond-XL nylon membranes (Cytiva) and hybridized to a radiolabeled 5′-(CCCTAA)5−3′ telomeric oligonucleotide probe. Radioactive signal imaging was performed with a Typhoon FLA 9000 imager (GE Healthcare) and quantified using ImageJ software.

### TERRA quantification

TERRA levels were quantified by RT-qPCR as described in (57, 58) with some modifications. Briefly, total RNA was extracted from the cells grown in 10 cm dish using QuickRNA Miniprep kit (Zymo Research) according to manufacturer’s protocol. For genomic DNA contamination removal, on-column DNAse I (10 Units) treatment was performed twice for 40 mins at room temperature, followed by in-solution DNase I treatment (8 Units, New England Biolabs) according to manufacturer’s instructions for 30 mins, at 37^0^C. RNA was subsequently re-purified using QuickRNA miniprep columns and subjected to a final DNAse I treatment (5 Units) on column for 30 mins at room temperature. For cDNA synthesis, 1 μg of RNA was reverse transcribed using TeloR (0.5 μM) and ActinR (0.05 μM) oligonucleotides (Table S9) and Superscript III (ThermoFisher Scientific) according to the manufacturer’s instructions. Quantitative PCR was performed to analyze TERRA RNA expression using the oligonucleotides listed in Table S9 and actin as a normalization control.

### Competitive proliferation assays

Nuclight lentivirus technology for live nuclear labeling (Sartorius) was used according to the manufacturer’s guidelines to distinguish sgCtl and SUB1 KO U2OS cells. Briefly, sgCtl and SUB1 KO cells were infected with either Nuclight Green (GFP fluorophore, Sartorius #4626) or Red (mKate2 fluorophore, Sartorius #4627) lentiviruses respectively, before being selected via bleomycin. Each cell line was then infected with lentiviruses expressing Cas9 and either a LacZ sgRNA (control) or a BLM sgRNA. After selecting infected cells with puromycin, GFP-positive and mKate2-positive cells infected by the same sgRNAs were seeded together at a 1:1 ratio. Mixed cells were grown over 20 days and cells were imaged daily using an automated fluorescent microscope (BD Biosciences, BD Pathway 855). GFP- and mKate2-positive cells were counted using ImageJ and statistical analyses were performed using GraphPad 10.

## Acknowledgments

We thank Rachel Litman-Flynn (Boston University) for sharing telomeric FISH protocols and cell lines and for critical reading of the manuscript. We thank Claus Azzalin, Bruno Silva and Sara Salgado (Instituto de Medicina Molecular João Lobo Antunes (Lisbon)) for sharing unpublished data and for fruitful discussions. We are indebted to Luc Gaudreau (U Sherbrooke) and Chantal Autexier (McGill) for the generous gift of cell lines and to Billy Gagnon for help with the cell line importation procedure. We thank Daniel Garneau for excellent support and training for fluorescence microscopy and image analysis. We are grateful to Manon Dufresne and Érika Milot for cell culture support. We thank all members of the Maréchal, Jacques and Wellinger labs for helpful discussions. Research in the Maréchal lab is supported by CIHR project grants (#376288 and #506553) and NSERC Discovery Grants (#4759) and PÉJ (#435710). Work in the Wellinger lab was supported by a CIHR Foundation Grant (#FDN154315) and the Canadian Research Chair on Telomere Biology. This research was enabled in part by support provided by Calcul Québec and the Digital Research Alliance of Canada (alliancecan.ca). This article is dedicated to the memory of Eliane B. Deschamps.

---

**Figure S1.**
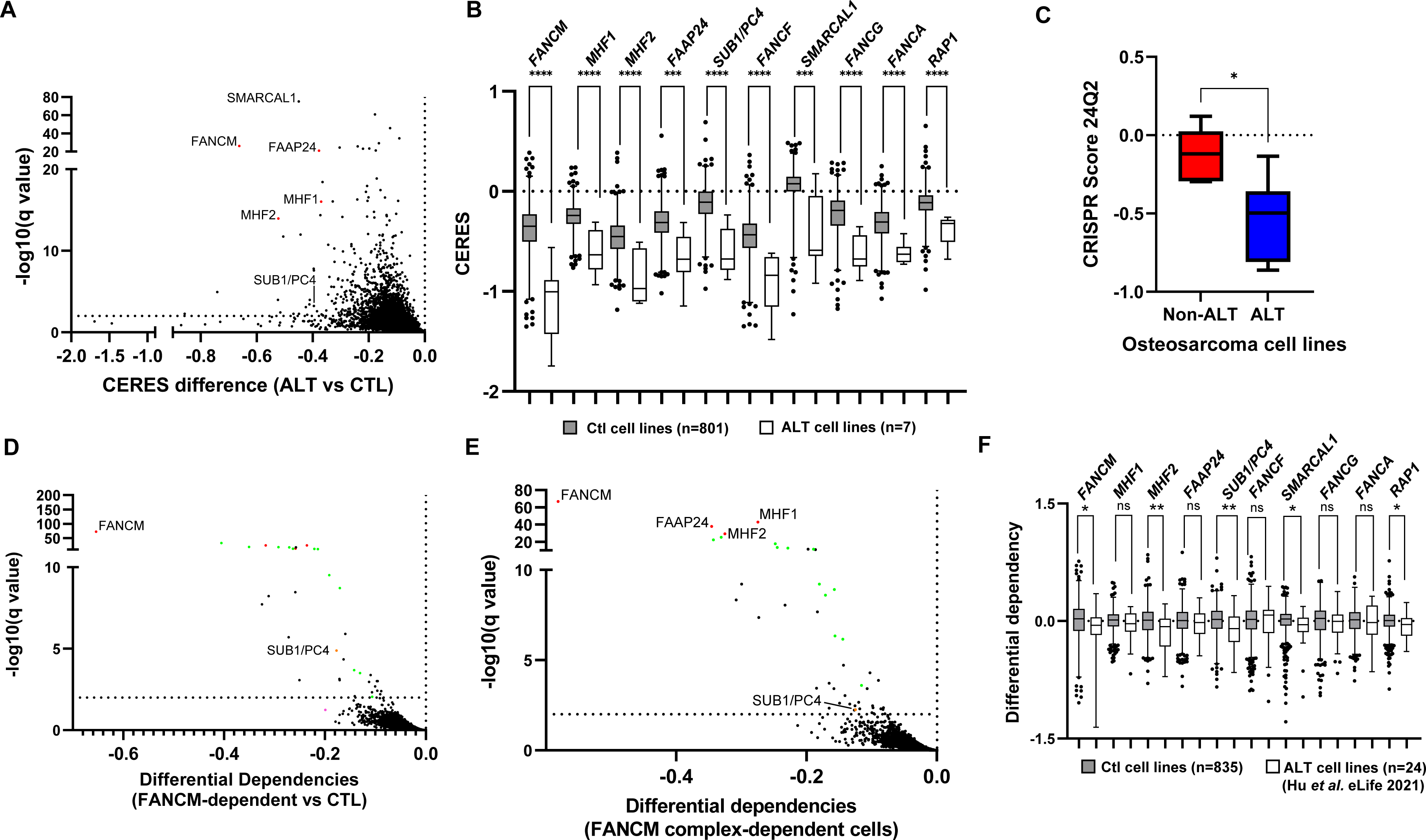
Identification of ALT-specific Vulnerabilities. **(A)** Volcano plot of the differences in CERES scores in experimentally validated ALT cell lines. CERES obtained from the Cancer Dependency Map (21Q1) for a panel of validated ALT cell lines (n=7) were compared with the rest of the available cell lines (n=801). CERES differences for ALT vs CTL cells were plotted against the FDR-corrected Q value, calculated from multiple unpaired t tests. The dashed horizontal line represents an FDR of 1%. **(B)** Box and whiskers plots of top hits in ALT cells. Horizontal lines represent the median and whiskers show 1-99 percentile cell lines. Significance was determined by Mann-Whitney tests. **** P<0.0001, *** P<0.001, ** P<0.01, * P<0.05. **(C)** The average SUB1 DEPMAP CRISPR scores (24Q2) of osteosarcoma cell lines with validated (ALT = 6) and non-validated (non-ALT = 5) ALT status were plotted along with standard deviations. Significance (P<0.05) was determined using Student’s t-test. **(D)** Volcano plot of the differential genetic dependencies in the top 5 % FANCM-dependent cells. Mean subtracted CERES scores obtained from the Cancer Dependency Map (21Q1) for a panel of FANCM-dependent cell lines (n=47) were compared with the rest of the available cell lines (n=761). Differential dependencies are plotted against the FDR-corrected Q value, calculated from multiple unpaired t-tests. The dashed horizontal line represents an FDR of 1%. Red dots represent FANCM complex proteins, green dots are Fanconi anemia factors. **(E)** Volcano plot of the differential genetic dependencies in the top 5 % FANCM-complex (FANCM/MHF1/MHF2/FAAP24) differentially dependent cells. Mean subtracted CERES scores obtained from the Cancer Dependency Map (21Q1) for a panel of FANCM-dependent cell lines (n=40) were compared with the rest of the available cell lines (n=768). Differential dependencies are plotted against the FDR-corrected Q value, calculated from multiple unpaired t-tests. The dashed horizontal line represents an FDR of 1%. Red dots represent FANCM complex proteins, green dots are Fanconi anemia factors. **(F)** Box and whiskers plots of preferentially essential genes of ALT cancer cell lines identified in Hu *et al.* eLife 2021 for which Depmap data was available (n=24) compared to control cell lines (n = 835) (21Q2). Horizontal lines represent the median and whiskers show 1-99 percentile cell lines. Significance was determined by Mann-Whitney tests (P<0.05 *, P<0.01 **).

**Figure S2.**
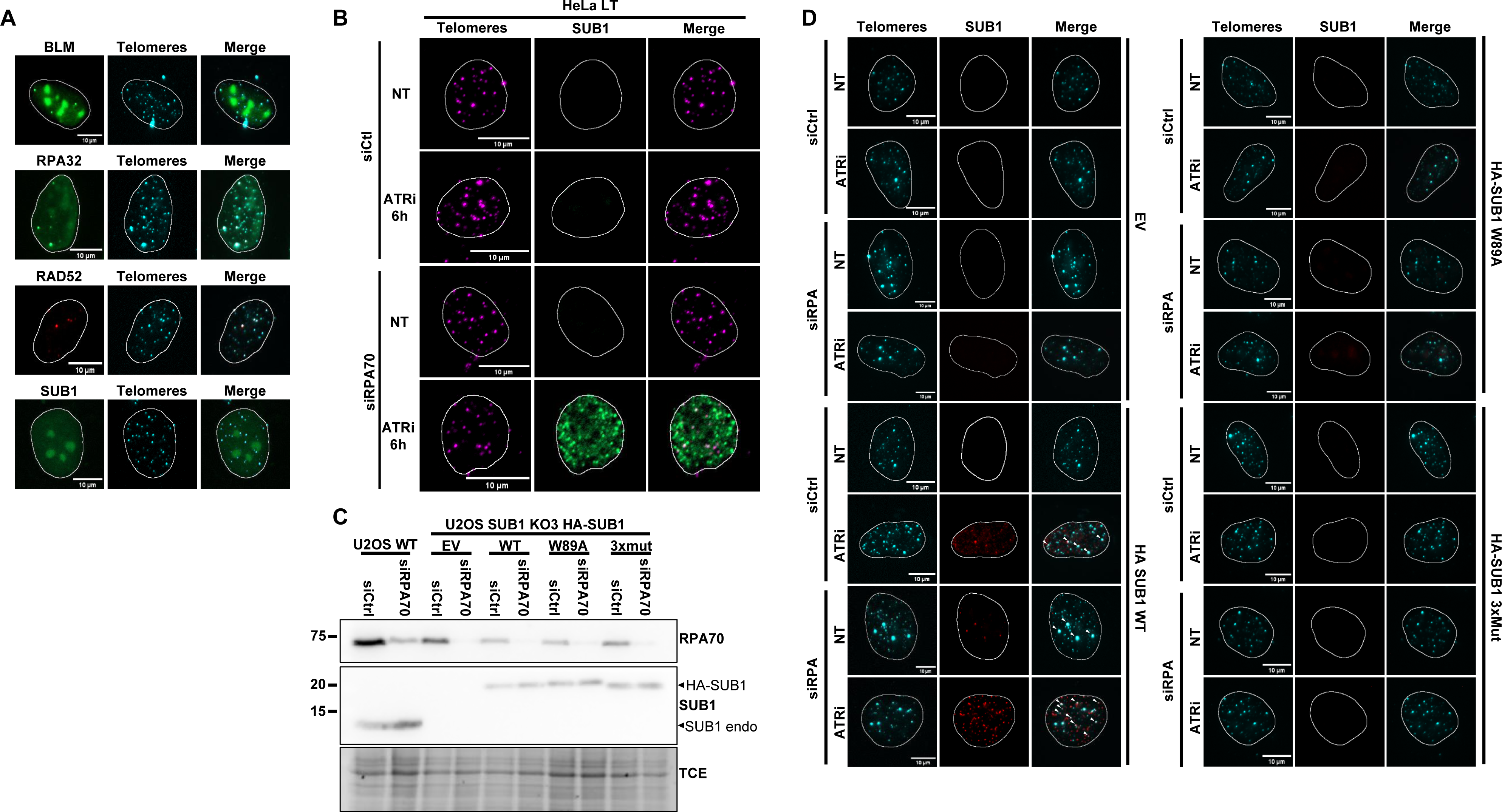
SUB1 localizes to damaged telomeres. **(A-C)** U2OS cells were processed for immunofluorescence and telomeric FISH with the indicated antibodies. **(A)** Representative images of BLM, RPA32, RAD52 and SUB1 telomeric foci in non-treated cells **(B)** Representative images of SUB1 WT foci in HeLa LT cells. **(C)** Immunoblot validation of RPA70 KD in WT and KO SUB1 cells complemented with the indicated HA-SUB1 constructs. **(D)** Representative U2OS KO SUB1 cells complemented with the indicated HA-SUB1 constructs or empty vector (EV) are shown. White arrows show colocalization events

**Figure S3.**
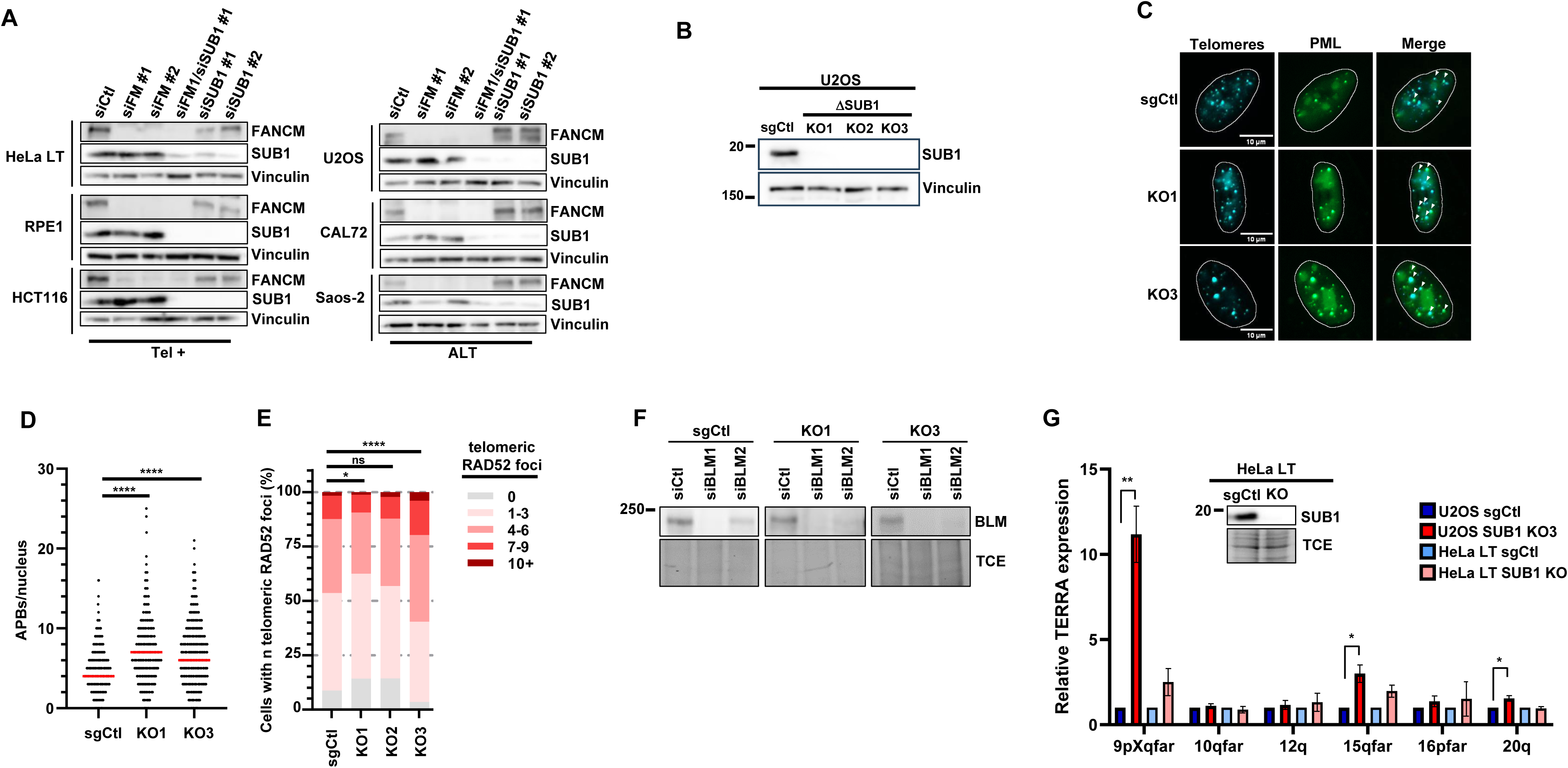
SUB1 alleviates RS at ALT telomeres. **(A)** Immunoblot validation of SUB1 and FANCM depletion in a panel of ALT and TEL+ cell lines. **(B)** Immunoblot validation of SUB1 KO U2OS cell lines. **(C,D,E)** sgCtl and KO SUB1 U2OS cell lines were processed for TEL-FISH, PML or RAD52 IF. ALT-associated PML bodies (APBs) and telomeric RAD52 foci were automatically counted using Cell Profiler. Experiments were performed in biological triplicates. Statistical significance was established by Kruskal-Wallis and Dunn’s multiple comparison tests (P<0.05 *, P<0.01 **, P<0.001 *** and P<0.0001 ****). **(F)** Immunoblot validation of BLM depletion in sgCtl and SUB1 KO cell lines. **(G)** RNA was extracted from sgCtl or SUB1 KO U2OS or HeLa LT cell lines and RT-qPCR was performed to amplify TERRA from the indicated chromosome ends. Data represents the mean relative TERRA expression from 3 independent biological replicates normalized to the sgCtl TERRA levels for each cell line. Statistical significance was determined using Student’s t-test (P<0.05 *, P<0.01 **).

**Figure S4.**
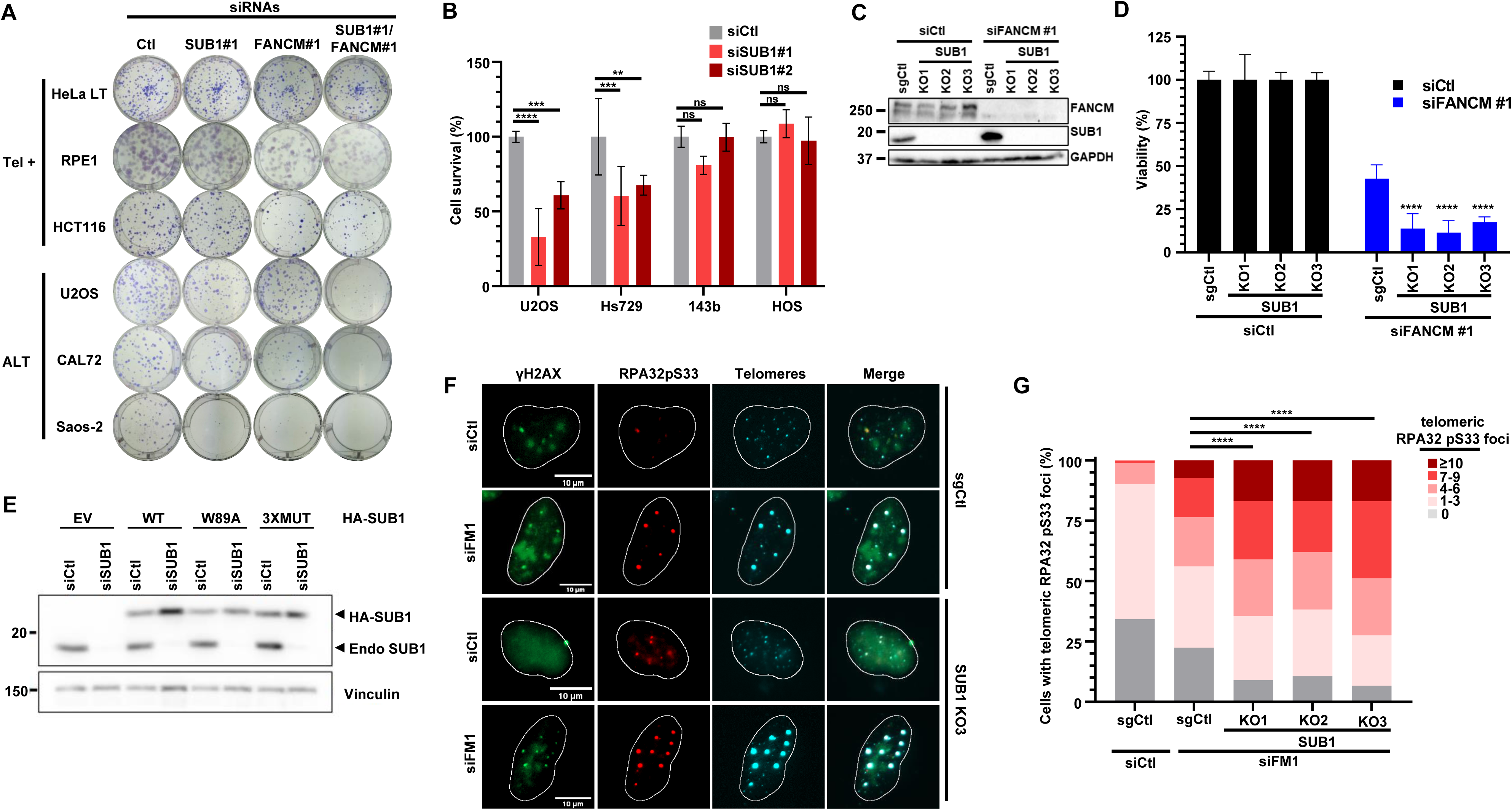
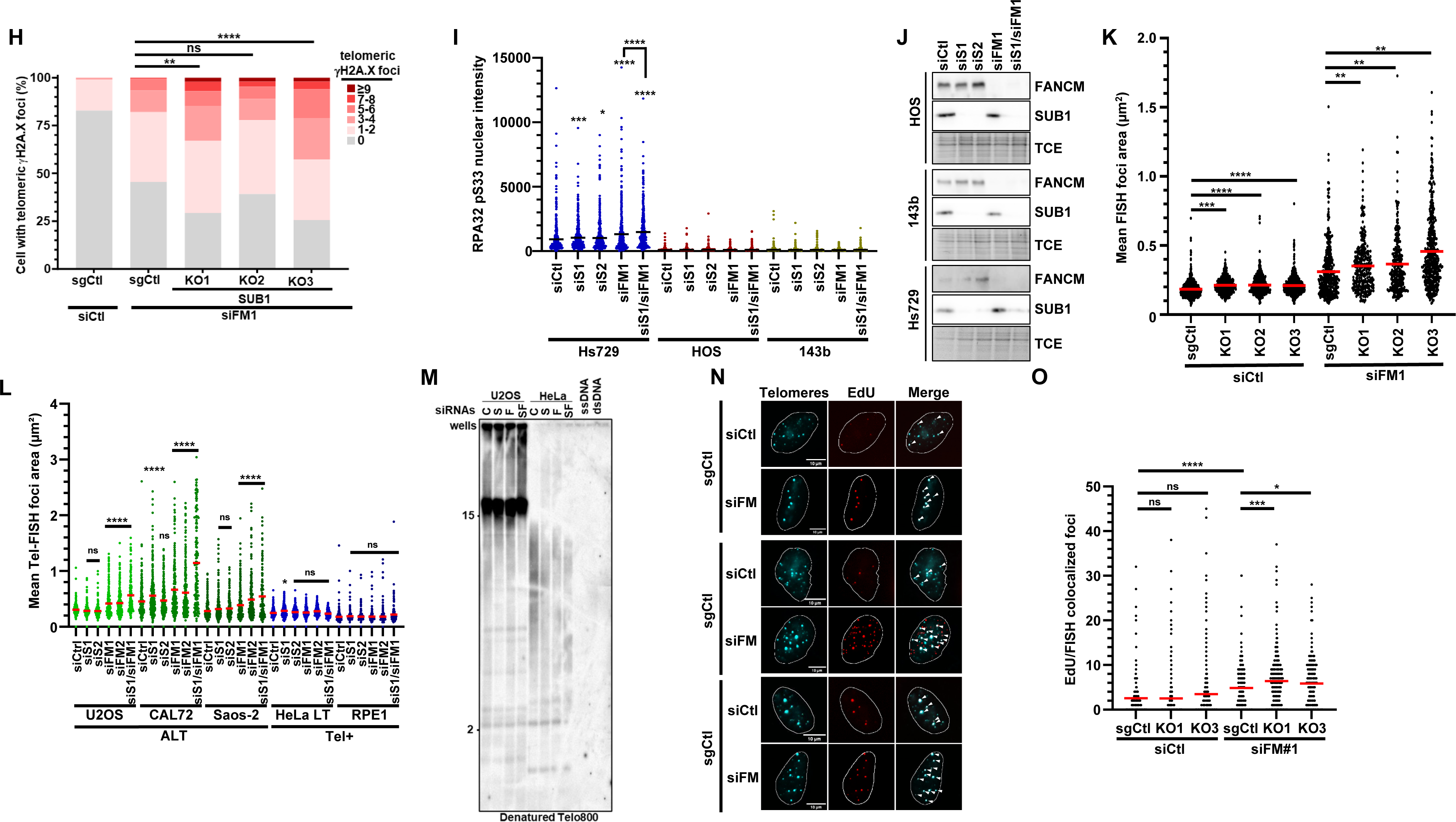
SUB1 and FANCM Co-Depletion is Deleterious for ALT Cells. **(A)** U2OS, CAL72, Saos-2, HeLa LT or RPE1 cells were transfected with the indicated siRNAs. 500 cells per condition were seeded onto 6-well plates and colonies were counted after 10-14 days. Representative images of colonies are shown. Viability was calculated compared to siCtl wells. **(B)** Clonogenic assays were performed for the indicated cell lines as in **A (C,D)** Immunoblot validation of SUB1 KO and FANCM depletion by siRNA in clonogenic assays **(E)** Immunoblot validation of endogenous SUB1 KD efficiency for colony formation assays. **(F,G,H)** SgCtl and KO SUB1 cells were transfected with the indicated siRNAs and IF for RPA32pS33 and γ-H2A.X along with telomere FISH were performed. Graph represents the merged data from three biological replicates for RPA32pS33 and two for γ-H2A.X. Statistical significance was established by one-way ANOVA and Šidák’s multiple comparison tests (P<0.05 *, P<0.01 **, P<0.001 *** and P<0,0001****). **(I)** Hs729, HOS and 143b cells were transfected with the indicated siRNAs and IF for RPA32pS33 was performed and nuclear intensity was quantified. Graph represents the merged data from biological replicates Statistical significance was established by one-way ANOVA and Šidák’s multiple comparison tests (P<0.05 *, P<0.001 *** and P<0,0001****). (**J**) Immunoblot validation of SUB1 depletion in the ALT rhabdomyosarcoma cell line HS729 and 2 independent non-ALT osteosarcoma cell lines. **(K)** Scatterplot of individual telomere foci area from sgCtl and KO SUB1 U2OS cells automatically quantified using Cell profiler. Each dot represents the mean telomere foci area/nuclei in µm^2^, red lines indicate the average. Graph represents the merged data from three biological replicates. **(L)** Mean telomere foci area from U2OS, CAL72 and Saos-2, HeLa LT and RPE1 transfected with indicated siRNA. Telomeric area was automatically quantified using Cell profiler. Graph represents the merged data from two biological replicates. **(M)** Normalization control for the native in-gel telomere hybridization experiment presented in 4E. After signal acquisition, gel was denatured and rehybridized with a radiolabeled long telomeric probe (Telo800). **(N,O)** U2OS sgCtl and SUB1 KO cells depleted using indicated siRNA were labelled with EdU and Click-it chemistry/IF were performed to visualize telomeric DNA synthesis. **(N)** Representative images of telomeric DNA synthesis **(O)** Graphical representation of the number of telomeric DNA synthesis instance per nuclei. Graphs represent the merged data from two biological replicates. Statistical significance was established by one-way ANOVA and Šidák’s multiple comparison tests (P<0.05 *, P<0.01 **, P<0.001 *** and P<0,0001****).

**Figure S5.**
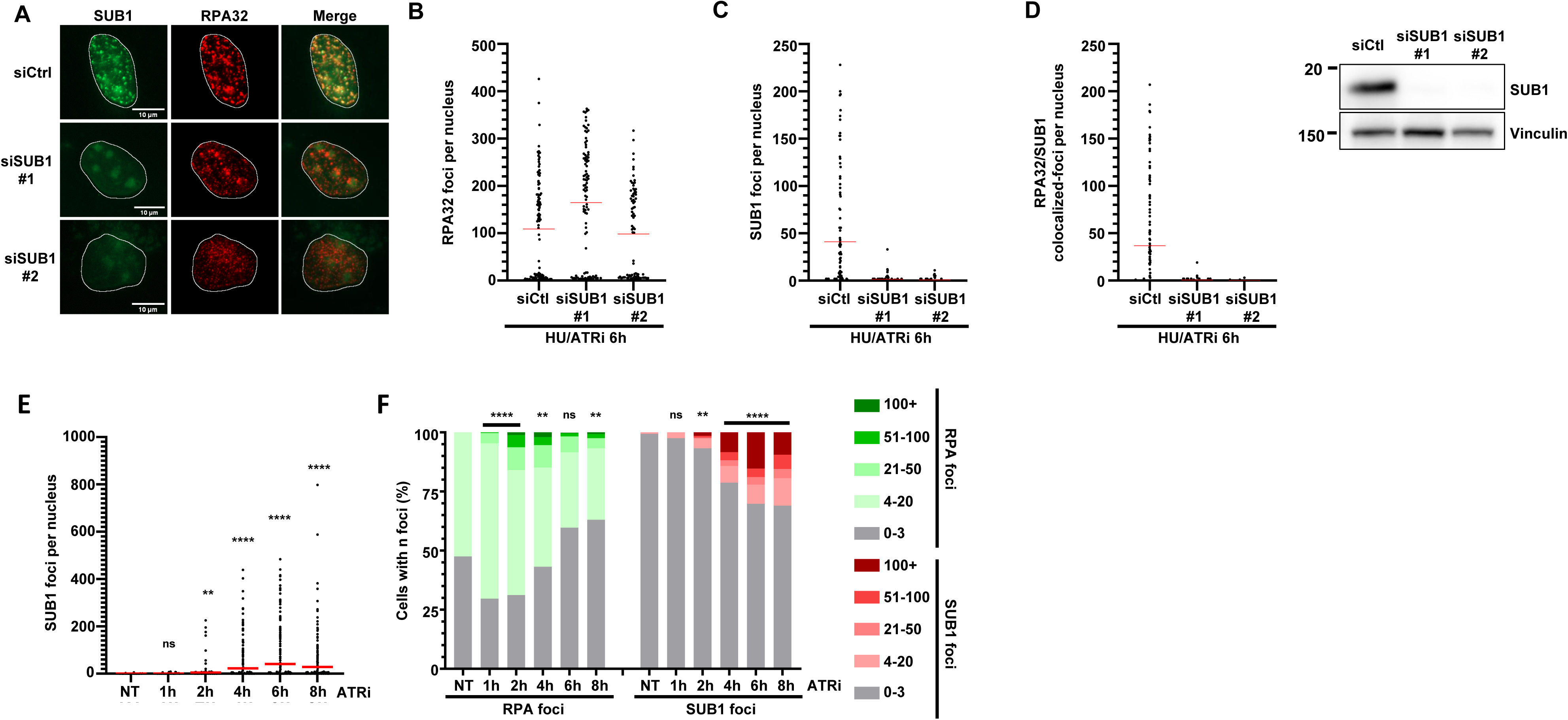
The single-stranded DNA binding protein SUB1 associates with stalled and collapsed DNA replication forks. **(A-D)** Specificity controls for SUB1 IF antibody. U2OS cells transfected with the indicated siRNAs were exposed to HU (2 mM) and ATRi (1 uM) for 6 hrs and processed for immunofluorescence. RPA32, SUB1 and colocalizing foci were automatically counted using CellProfiler. **(E,F)** U2OS cells were treated with ATRi (10 uM AZD6738) for the indicated times and processed for immunofluorescence against SUB1 and RPA32. **(E)** Graphical representation of the number of SUB1 foci in each nucleus. **(F)** Histogram of the proportion of ATRi-treated cells with 0-3, 4-20, 21-50, 51-100 or 100+ RPA32- and SUB1-foci. Graph represents a single replicate and experiments were performed in biological replicates with similar results. Statistical significance was established by one-way ANOVA and Dunn’s multiple comparison tests (P<0.01 **, P<0.0001 ****).

**Figure S6.**
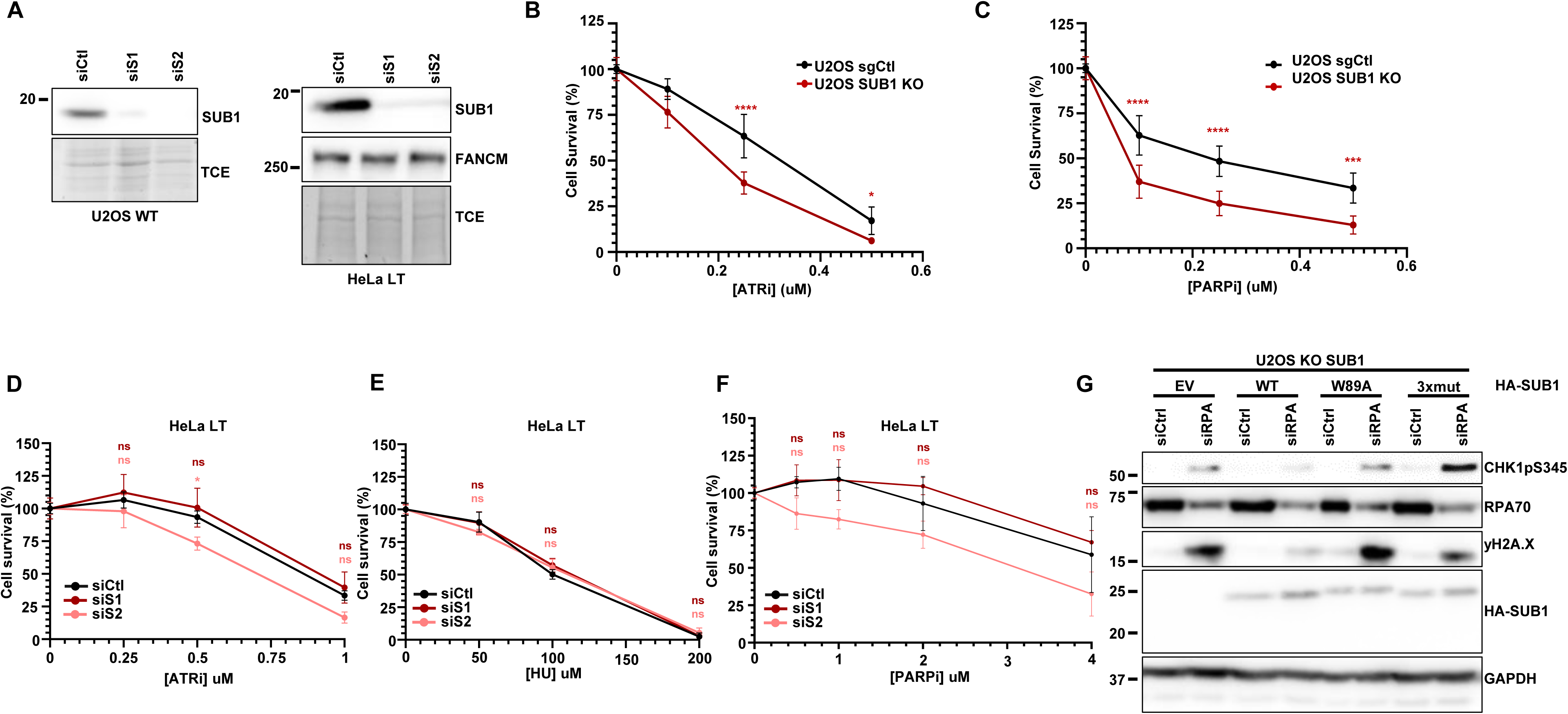
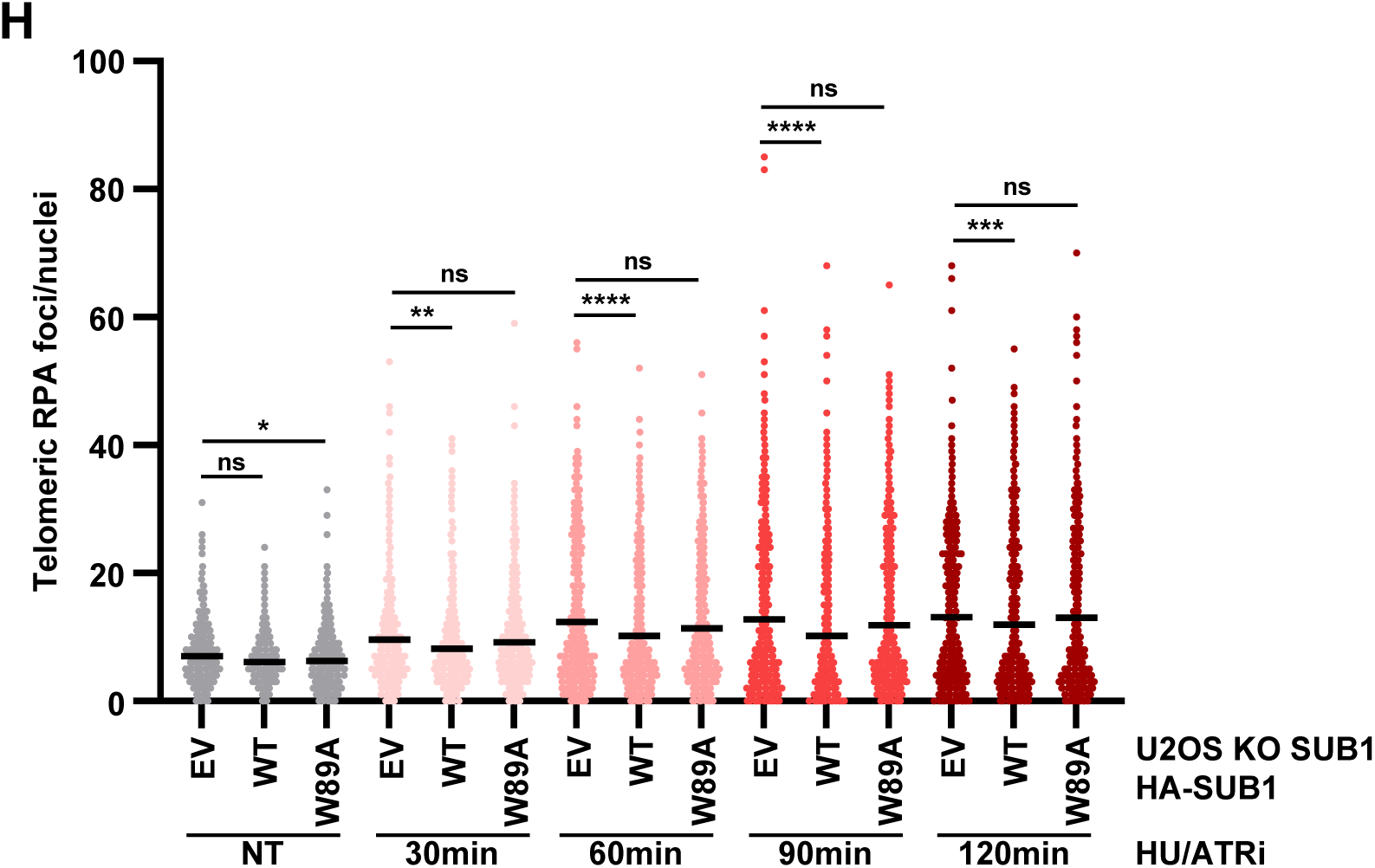
SUB1 Protects Against RS and Replication Catastrophe in ALT cells. **(A)** Immunoblot validation of SUB1 depletion in U2OS and HeLa LT cells. **(B,C)** sgCtl or SUB1 KO U2OS cells were treated with the indicated drug concentrations for 4 days and allowed to form colonies over 10 days prior to fixation and viability assessment. Graphs represent the merged data from three biological replicates **(D,E,F)** HeLa LT cells were transfected with the indicated siRNAs and incubated with HU, ATRi or PARP1i at the indicated concentrations as done above. Graphs represent cell viability compared to non-treated cells. Graphs represent the merged data from three biological replicates each consisting of two technical replicates. Statistical significance was established by ordinary two-way ANOVA and Tukey’s multiple comparison tests. **(G)** WT and KO SUB1 cells complemented with the indicated HA-SUB1 constructs were transfected with the indicated siRNAs and 48 hrs later lysed and processed for immunoblotting using the indicated antibodies. **(H)** U2OS SUB1 KO3 cells complemented with an empty vector HA-SUB1 WT or HA-SUB1 W89A were treated with 2mM HU and 1μM ATRi for indicated amount of time and processed to visualize Telomeric RPA by IF (RPA70/TRF2). TRF2 and RPA foci were automatically counted using CellProfiler. Statistical significance was established by Kruskal-Wallis and Dunn’s multiple comparison tests (P<0.05 *, P<0.01 **, P<0.001 *** and P<0.0001 ****).

**Figure S7.**
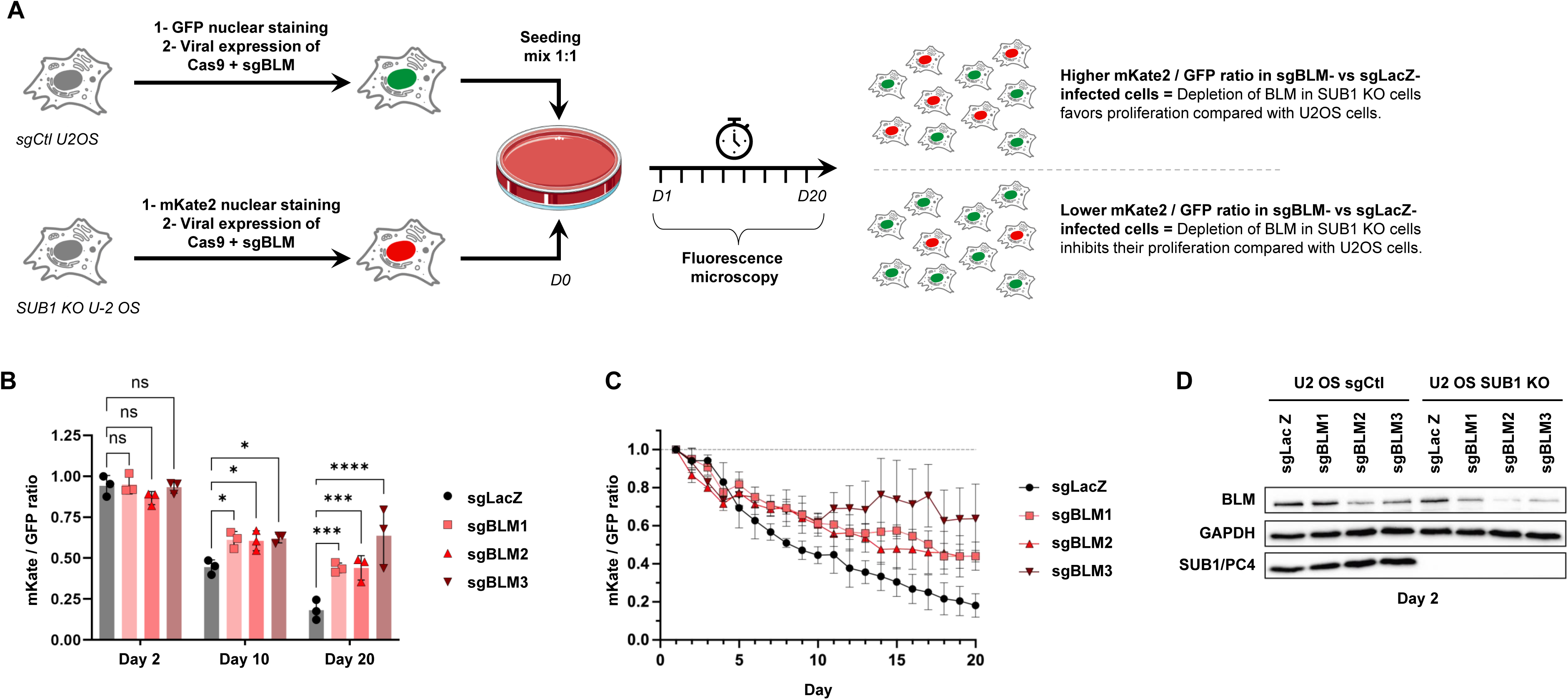
BLM depletion stimulates the proliferation of SUB1 KO cells. **(A)** Competitive assay workflow. Nuclei of sgCtl and SUB1 KO U2OS cells were labelled using GFP and mKate2 fluorophores respectively. Both cell lines were infected with lentiviruses carrying Cas9 and sgRNAs targeting LacZ or BLM. Puromycin-selected cells were mixed at a 1:1 ratio. The mKate2/GFP cell ratio was assessed daily over 20 days using fluorescence microscopy. As SUB1 KO U2OS (mKate2) cells grow slower than control U2OS cells (see decreasing mKate2/GFP ratio over time in sgLacZ-infected cells), a higher mKate2/GFP ratio over time in sgBLM-infected cells compared to sgLacZ-infected ones indicates that BLM depletion promotes the growth of SUB1 KO cells compared with sgCtl U2OS cells. **(B)** Growth competition assay between sgCtl and SUB1 KO U2OS cells, following viral knockdown of BLM. sgCtl U2OS were labelled using GFP while SUB1 KO U2OS cells were identified using mKate2. Plotted values represent the ratio of the mKate2+ cells versus the GFP+ cells; normalized to day 1. Statistical significance was established by ordinary two-way ANOVA and Dunnett’s multiple comparison tests (p-values: P<0.05 (*), P<0.001 (***) and P<0.0001 (****)). **(C)** Time course of the growth competition assay. **(D**) Immunoblot validation of BLM depletion in the sgCtl and SUB1 KO U2OS cells at day 2.

## Notes

### Competing Interest Statement

The authors have declared no competing interest.

